# Estimation of spatial demographic maps from polymorphism data using a neural network

**DOI:** 10.1101/2024.03.15.585300

**Authors:** Chris C. R. Smith, Gilia Patterson, Peter L. Ralph, Andrew D. Kern

**Affiliations:** Institute of Ecology and Evolution, University of Oregon, Eugene, OR 97403, USA

## Abstract

A fundamental goal in population genetics is to understand how variation is arrayed over natural landscapes. From first principles we know that common features such as heterogeneous population densities and barriers to dispersal should shape genetic variation over space, however there are few tools currently available that can deal with these ubiquitous complexities. Geographically referenced single nucleotide polymorphism (SNP) data are increasingly accessible, presenting an opportunity to study genetic variation across geographic space in myriad species. We present a new inference method that uses geo-referenced SNPs and a deep neural network to estimate spatially heterogeneous maps of population density and dispersal rate. Our neural network trains on simulated input and output pairings, where the input consists of genotypes and sampling locations generated from a continuous space population genetic simulator, and the output is a map of the true demographic parameters. We benchmark our tool against existing methods and discuss qualitative differences between the different approaches; in particular, our program is unique because it infers the magnitude of both dispersal and density as well as their variation over the landscape, and it does so using SNP data. Similar methods are constrained to estimating relative migration rates, or require identity by descent blocks as input. We applied our tool to empirical data from North American grey wolves, for which it estimated mostly reasonable demographic parameters, but was affected by incomplete spatial sampling. Genetic based methods like ours complement other, direct methods for estimating past and present demography, and we believe will serve as valuable tools for applications in conservation, ecology, and evolutionary biology. An open source software package implementing our method is available from https://github.com/kr-colab/mapNN.

## 1 Introduction

To study, conserve, and manage populations, we must characterize where individuals live and how they move. For instance, differences in habitat suitability cause local geographic regions to support higher or lower numbers of individuals. Similarly, landscape features such as mountain ranges restrict dispersal to or from different areas. Understanding local population density and dispersal patterns is important for prioritizing conservation resources, for predicting species range shifts in response to ongoing climate change, and for studying hybrid zones and speciation. Here, we describe a novel approach for estimating these ecologically important parameters from single nucleotide polymorphism (SNP) data, a form of genetic data readily available from many non-model species.

First, we give a brief overview of available methods to highlight the need for further development of molecular tools. It is often not feasible to count every individual in a population and in these cases researchers rely on statistical inference to estimate population sizes. Perhaps the most commonly used approach, mark-recapture, is sampling intensive: thousands of captured individuals may be required to analyze large populations (Schorr et al., 2014). Likewise, it is rare to have access to complete or even near-complete dispersal information for a population, although something close to this might be achieved with radio tracking collars, e.g., with wolves. As an alternative, population genetic variation can provide information about density and dispersal using relatively small sample sizes, although this information is less direct and often reflects averages over longer time scales. This option becomes even more attractive as technologies for DNA sequencing and non-invasive genetic sampling continually improve.

While estimates of census population sizes are often the goal for ecological or conservation applications, population genetics methods instead typically infer *effective* size, *N_e_*. Indeed numerous sophisticated methods have been developed for this task. For example, PSMC (Li and Durbin, 2011) estimates past changes of *N _e_* though time from a single diploid genome using coalescent theory combined with a hidden Markov model. However, PSMC and related methods assume the studied population is well-mixed and mating randomly, whereas real populations are often highly structured, which leads to strong differences between *N_e_* and the census size of a population. For example, Battey et al. (2020b) showed using simulations that genetic diversity varied by almost a factor of four between well-mixed populations and those with significant spatial structure due to limited dispersal. This effect will be increased for species with large ranges relative to dispersal rate, and suggests that genetics-based population size inference will benefit from explicitly including geography.

Population genetics has also been used to estimate *spatial* demographic parameters. The method of Rousset (1997) uses the slope of the least squares fit of genetic distance versus geographic distance to estimate “neighborhood size”: the mean number of nearby, potential mates. However, neighborhood size is a function of both the effective population density, *D*, and effective dispersal rate, *σ*, and it is challenging to disentangle the two parameters if both are unknown. Even so, if either value is known for a population, the other can be estimated using Rousset’s formula. More recently, we presented a neural network trained on simulated data that estimates *σ* when *D* is unknown (Smith et al., 2023). Ringbauer et al. (2017) developed a related approach for estimating both *D* and *σ* together using identity-by-decent blocks as input. Although useful, the Rousset (1997) and Ringbauer et al. (2017) methods, as well as our original method, assume a homogeneous environment.

While the aforementioned studies focused on estimating a single population-wide density and dispersal rate, these quantities likely vary across the landscape. Such spatial heterogeneity will not only bias our population-wide inferences but represents an opportunity to learn even more from geo-referenced SNP data. There are of course other tools to explore spatially varying demography. The most commonly used are `EEMS` (Petkova et al., 2016) and its updated version, `FEEMS` (Marcus et al., 2021), which model obstacles to gene flow using an analogy to electrical resistance (McRae, 2006; McRae et al., 2008) and are designed to estimate effective migration surfaces. These programs have been widely used for characterizing population structure and landscape connectivity. However, the effective migration surfaces from `EEMS` and `FEEMS` do not provide separate estimates for population density and dispersal rate. Furthermore, these (and other resistance distance-based methods) approximate the coalescing of two lineages as a single commute on a graph. In practice, these methods can be strongly affected by realistic scenarios like biased migration (Lundgren and Ralph, 2019) or density variation. `Circuitscape` (McRae et al., 2009) is a related tool for obtaining connectivity maps from genetic data, however it is distinct from the above methods because it uses the additional input of known or inferred landscape layers. However, in the current study we focus on methods that do not require landscape features or environmental data. To address the above limitations, Al-Asadi et al. (2019) created `MAPS` which uses identity by descent input to estimate population density and dispersal rate, and does not depend on the resistance distance approximation. At present however, high quality genomic resources such as identity by descent blocks remain out of reach for most non-model organisms, thus it is important to develop new methods that are compatible with more accessible input data.

A promising avenue for dealing with high-dimensional, geo-referenced genotypes is to train a deep neural network to identify useful features in the data in an automated fashion (Smith et al., 2023). Deep neural networks can be trained on simulated data, which bypasses the need to obtain empirical data for training. Such models in some cases outperform traditional methods in population genetics, and hold the potential for innovative analyses that address longstanding questions. They are particularly well-suited for inference with heterogeneous, biologically realistic quantities such as maps, where analytical expectations are difficult to obtain. In recent years, supervised deep learning approaches have been developed for various tasks in population genetics, for example: inferring demographic history (Sheehan and Song, 2016), detection of selective sweeps (Kern and Schrider, 2018), detecting introgression (Ray et al., 2023), identifying geographic origin of an individual using their DNA (Battey et al., 2020a), estimating mutation rate (Fang et al., 2022), estimating recombination rate (Adrion et al., 2020), and characterizing admixture (Oriol Sabat et al., 2022). In the realm of spatial analysis, our method `disperseNN` used a deep neural network trained on simulated SNP data to infer dispersal rate (Smith et al., 2023). A recently updated model (Smith and Kern, 2023) implemented a pairwise convolutional network, which was particularly effective at estimating the spatial demographic parameter of interest.

Here, we present a new method, `mapNN`, for estimating spatial maps of demographic parameters from polymorphism data. The approach requires custom, simulated datasets for training, and provides estimates for population density and dispersal rate across the landscape. `mapNN` was designed to work with SNPs and relatively small sample sizes, making it well suited for non-model species. The software is available from: https://github.com/kr-colab/mapNN. In addition to describing the new method, we benchmark `mapNN` against existing methods for estimating demographic maps, and apply it to empirical data from North American grey wolves.

## 2 Materials and Methods

### 2.1 Simulations

The `mapNN` approach relies on simulations for both training and testing. The following procedure was used to generate the simulated data.

#### 2.1.1 Continuous space population model

Simulations were carried out using the continuous space, individual-based population genetic simulator, `SLiM` (Haller and Messer, 2023). Our `SLiM` model is similar to those of Battey et al. (2020b) and Smith et al. (2023). Each life cycle consists of reproduction, dispersal, and mortality. Individuals are hermaphroditic, and mates for a given individual are chosen with probability proportional to a Gaussian density about that individual with mean zero and standard deviation *σ_m_*, out to a maximum distance of 3*σ_m_*. The number of offspring produced per mating is Poisson distributed with mean 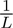, where *L*, the (rough) average lifetime at stationary, is 4. Dispersal occurs as new offspring are initially placed onto the habitat map: dispersal from the maternal parent is Gaussian distributed with mean zero and standard deviation *σ_f_* in both the *x* and *y* dimensions. Offspring that would land outside the habitat boundary are not produced. Mortality is competition based, where the probability of survival for individual *i* is 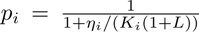; here, *K _i_* is the carrying capacity at the individual’s location, and *η_i_*is the sum of competitive interactions of the individual with their neighbors, which is calculated using a Gaussian kernel with standard deviation *σ_c_* up to a maximum distance of 3*σ_c_*. `SLiM` does not assume a particular unit of spatial distance.

The dispersal value we intend to estimate is *σ*—the root mean square distance along a spatial axis to a random parent—, which is an outcome of the simulation rather than an input parameter. In our simulation model, *σ* is modulated primarily by the *σ_f_* and *σ_m_*parameters. In Smith et al. (2023) we showed that *σ_c_* has a negligible effect on effective dispersal. The other value we estimate is population density, *D* —the number of individuals per unit area—, which is determined mainly by local carrying capacity.

To incorporate spatial heterogeneity we define a *w* × *w* spatial grid, where grid cells can carry different values for carrying capacity and dispersal rate. We then use `SLiM`’s built-in functionalities to read an input file of comma separated values describing the dispersal rate and carrying capacity values at each grid cell in the map, where continuous values for individuals are bilinearly interpolated.

#### 2.1.2 Generating demographic maps

Our strategy for creating input maps was to generate random spatial segments, where each segment has a different value for carrying capacity and dispersal rate (Figure 1). The minimum number of segments in our maps was one, representing a “flat” map. The maximum number of segments among maps is a tunable hyperparameter that would likely affect the spatial precision of the inferred maps, although we have not experimented with this. We used a maximum number of segments of three.

**Figure 1:**
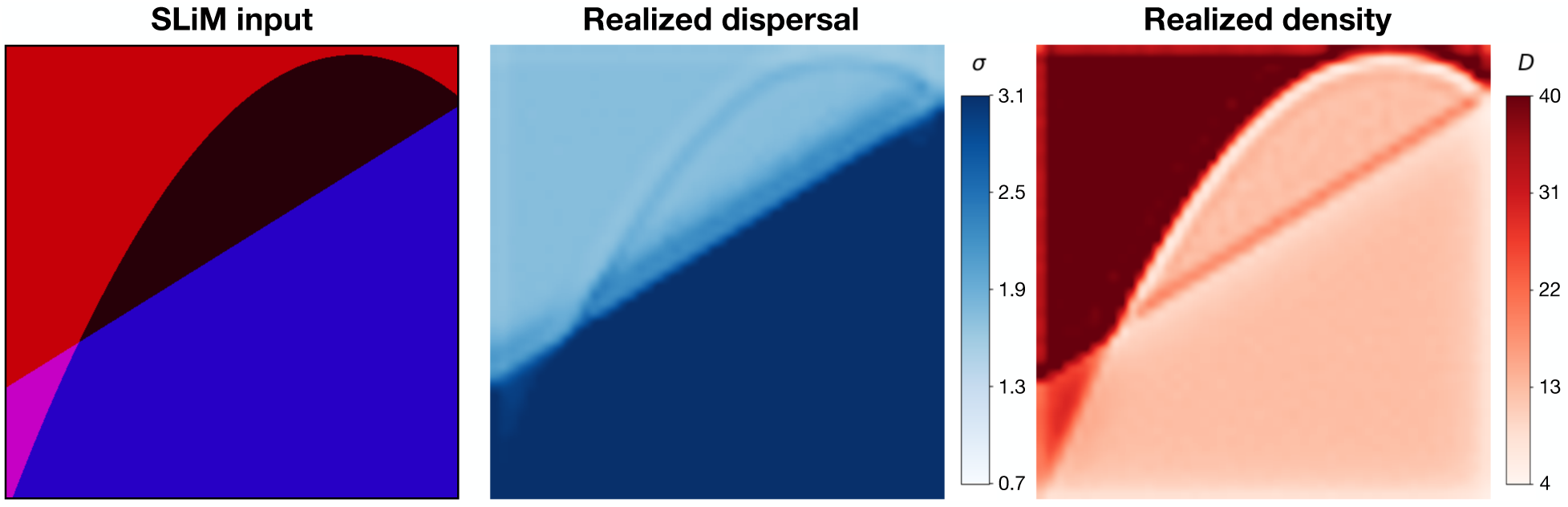
Example training map. The leftmost panel is a PNG rendering of the input map for `SLiM` : the magnitude of red conveys local carrying capacity for each cell in the map grid and the blue channel conveys dispersal rate. The middle panel is a heat map of the realized dispersal rate tracked during the simulation, using a 50×50 grid and other parameters as described in the main text. The right-hand panel is a heat map of realized density tracked during the simulation.

To create the boundaries between the distinct segments of the map, we first draw a random degree for a polynomial curve uniformly between zero and three (inclusive). If the sampled degree is zero, we draw a flat map. Otherwise, we draw a polynomial function of the chosen degree, i.e., a straight line, a quadratic curve, or a cubic. A number of random points equal to the degree plus one are uniformly sampled from the landscape, and the polynomial curve is fit to the chosen points. For instance, if the degree is two, then the boundary between the regions will be the curve *y* = *ax* ^2^ + *bx* + *c*, where *x* and *y* are the coordinates of the map and *a*, *b*, and *c* are chosen so the curve passes through three uniformly chosen points in the range. This splits the range into some number of regions. For the flat map case, we assign a log uniform value from a “prior” range (*p_min_, p_max_*) for the parameter of interest. For maps with multiple segments, we first draw a uniform value from (*p_min_, p _max_*)—this will become the range of values on the map, *r*. Next, we draw a log-uniform value from (*p_min_, p_max_* − *r*), to represent the minimum map value, *m*. If the number of segments is two, they are randomly assigned *m* and *m* + *r*. Otherwise, we draw additional values uniformly from (*m,m* + *r*) for the intermediate segments. We assign each of 50 × 50 grid cells to a segment based on whether the location of the cell falls below the curve. Last, we randomly flip and rotate each map.

Maps of dispersal and density are independently generated using the same procedure and then stacked together in a three dimensional array. See Figure S1 for a sample of training maps from the benchmark analysis.

#### 2.1.3 Recording population density and dispersal

We expected the realized population density and dispersal rates to differ somewhat from the values in the input map, due to random noise and also due to complex dynamics related to non-uniform demography. In order to obtain “true” maps of demography, we recorded local density and dispersal events throughout the benchmarking simulations (see below) in a 50 x 50 grid, excluding the first 250 generations to allow the spatial aspects of the simulation to reach a somewhat steady state (with continual fluctuation). For example, during the initial generations of each simulation the population size changed moderately from the initialized number of individuals to an equilibrium size. However, different burn-in times may be necessary for different simulation models or parameterizations. These “ground truth” maps were used for validating `mapNN` and the other inference methods.

The local population density and dispersal rate realized during the simulation closely resembled that of the *K* and *σ* channels in the input map (Figure 1). The correspondence between realized and expected *σ* values was quite good: the input *σ* was 1.9% smaller than the realized *σ* on average (SD=2.4%). However, the correspondence with expected density was imperfect: *K* was 14.0% larger than the actual population density (SD=2.6%). The exact relationship of the parameters *σ* and *K* to the realized average dispersal distance and density is somewhat complex, particularly when the parameters vary spatially. For instance, the edge of a high-dispersal area bordering a low-dispersal area has reduced density because there is a net flux away from the area. Similarly, a low density area bordering a high density area has a larger proportion of longer-distance migrants. However, the correspondence is fairly close in our examples.

#### 2.1.4 Sampling

In `mapNN` we have implemented three different strategies for sampling individuals from simulations. (1) Individuals may be sampled uniformly at random from those alive at the sampling time. It is important to note that if the environment is heterogeneous, uniform sampling will result in unbalanced sampling across space. (2) Alternatively, sampling may follow a grid where a set number of individuals are sampled from each grid cell. This approach can provide near-uniform sampling with respect to space, but it requires that all grid cells contain sufficient individuals. (3) We also provide the option to use a set of arbitrary, “fixed” locations. Here, we sample the individual that is nearest to each of the provided coordinates. The latter can be used to recapitulate an empirical sampling scheme in the training data.

#### 2.1.5 Generating neutral genetic variation

To initialize the simulations, we used genetic diversity simulated using the coalescent model implemented in `msprime` (Baumdicker et al., 2022). The coalescent portion is referred to as “recapitation” and was needed to reduce computation time. This two-step strategy—`SLiM`, then recapitate—means that recent generations are spatially explicit, while older generations approximate a Wright-Fisher model (i.e., with random mating). Mutations were simulated along the genealogy of the sampled individuals using `msprime`. Rather than use a fixed mutation rate, we simulate a fixed number of polymorphic sites; this is accomplished by initially adding mutations with a small rate (*µ* = 1 × 10^−15^), and then iteratively multiplying *µ* by a factor of two and simulating additional mutations until the desired number of SNPs, *m*, is found, at which point we uniformly sample *m* SNPs. This works well in practice, because maintaining a constant number of variable sites makes training more efficient by avoiding the need for zero-padding or variable-width tensors. In a validation setting, it challenges the neural network to estimate demographic parameters using information other than the number of segregating sites in a sample. And in empirical applications, it lets us match the number of SNPs in a focal, empirical dataset, which should maximize accuracy.

### 2.2 Neural network architecture

Our deep learning model (Figure 2) was developed using the Tensorflow (Abadi et al., 2016) and Keras (https://github.com/keras-team/keras) libraries. The model has two input branches, one that extracts features from genotypes and one that learns the relationship between genotype locations and grid cells in the map. The genotype-feature extraction branch uses a pairwise convolutional network similar to that of Smith and Kern (2023). This network loops over all pairs of individuals, sharing model parameters across pairs, and outputs a set of features for each pair. This strategy summarizes the relatedness between pairs of individuals and, as we have shown in Smith and Kern (2023), is effective at estimating the population-wide dispersal rate. The input to the first branch is a genotype matrix of 0s, 1s, and 2s representing the count of the minor allele for each SNP from each individual; although, `mapNN` also has an option for phased SNP data which would consist of two haplotypes per individual with 0s and 1s. After multiple iterations of 1-dimensional convolution and pooling layers, the intermediate features are flattened and put through a fully connected layer; the output of this layer is a set of 128 machine-learned genotype summaries for each pair of individuals. The above procedure is repeated for all 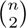 pairs of individuals (or a random subset of pairs; see below), and the features from each pair are concatenated together to form *G*, a matrix with one row per pair and 128 columns. The dimensions and activation function of each layer are described in Figure S2.

**Figure 2:**
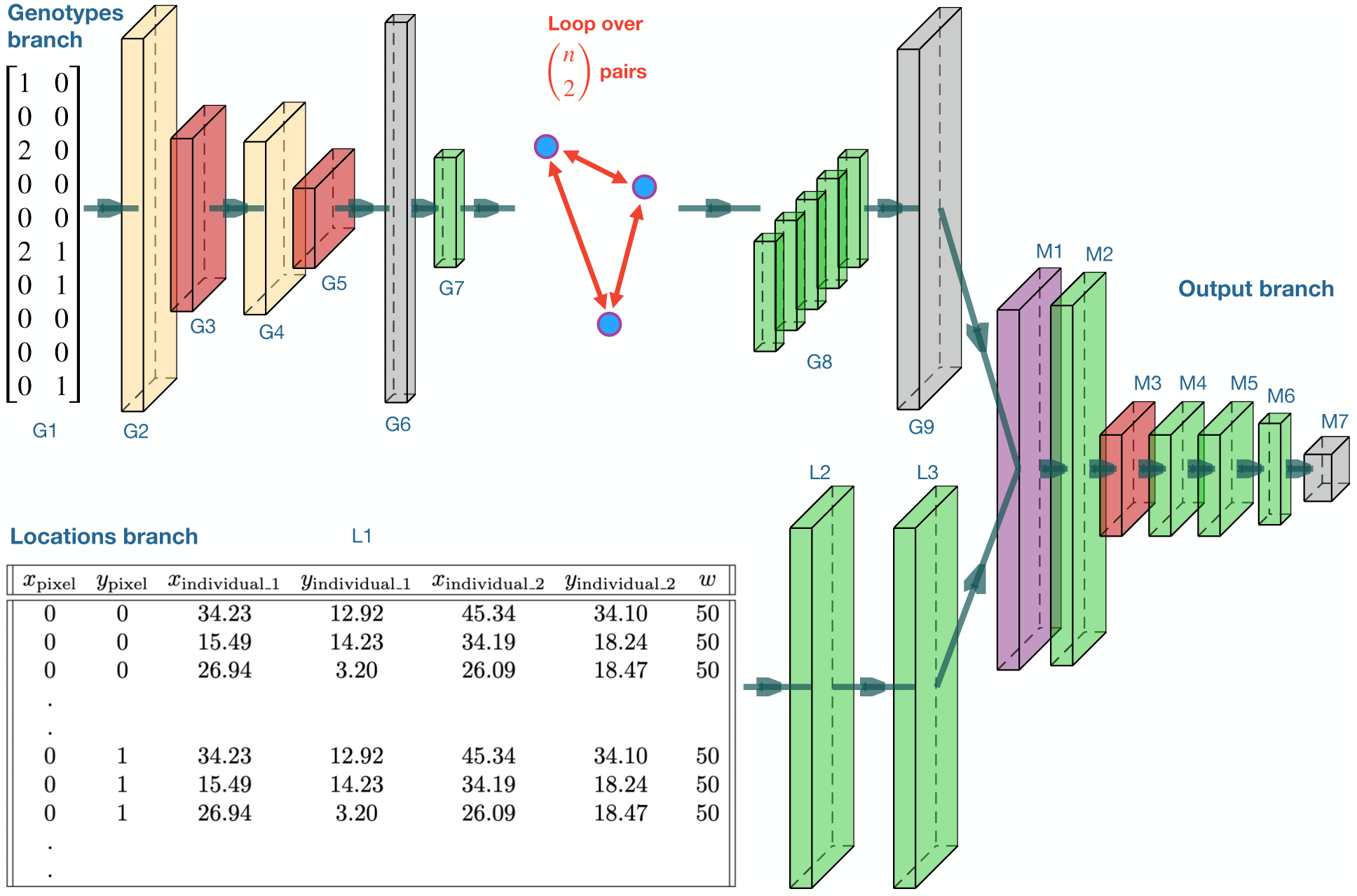
Neural network schematic for `mapNN`. The genotype branch loops over pairs of individuals— shown is a table of ten SNPs for an example pair of genotypes, but larger numbers of SNPs are used in the actual analysis. Rectangular prisms convey neural network layers, colored by type: convolution (beige); pooling (red); flattening, duplication, or reshaping (grey); fully connected (green); and elementwise multiplication (purple). Descriptions and output sizes for each layer are described in Figure S2. The example locations input table is truncated for visualization and is described in the main text. The output is a square map with separate channels for dispersal and density. This image was generated using https://github.com/HarisIqbal88/PlotNeuralNet.

We reasoned that to estimate the value of a grid cell in the map, it would be important to utilize genotype features from all pairs of individuals simultaneously, thus maximizing usage of the available information. However, the individuals in a pair are spatially closer to some grid cells than to others. Therefore, we sought to weight the genotype summaries 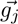 or the *j* th pair according to that pair’s spatial relevance to each cell. As an example, the spatial weighting could represent the mean Euclidean distance between cell *i* and the individuals in pair *j*. Of course, the Euclidean distance is only one of many ways to summarize the relationship between cells and pairs and thus might limit the behavior of the ultimate model. For example, grid cells along the line segment between two individuals would ideally carry information about genetic connectivity. As it is challenging to know the relevant spatial variables *a priori*, we tasked the neural network with determining the most useful spatial scores relating each pair to each grid cell in the map—this is the task of the second network branch.

For the second input, we put together a table containing one row of geographic coordinates for every combination of grid cell and pair (Figure 2; specifically, each row has the *x* and *y* coordinates for a cell, the coordinates of the first individual in a pair, the coordinates of the second individual in the pair, as well as the map width, *w*). This table contains redundant information, but is effective for conveying the spatial relationship between individuals and map locations to the network. The locations are processed using two dense layers, which leads to a number of spatial scores equal to the number of genotype features. By extracting multiple spatial scores we hope to produce a rich feature space that is used for estimating the value of each cell in the map grid. ReLU activation functions constrain spatial scores to be non-negative. This leads to a matrix, *S_i_*, for the *i* th cell containing a row of 128 spatial scores for each pair (Figure S2).

The final portion of the network applies the spatial weights to the genotype features, and estimates the grid values constituting the spatial map of demographic parameters. The values of a given cell in the map are calculated as:

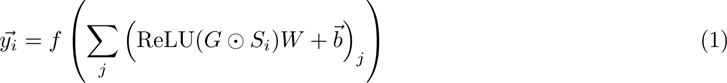

where ⊙ denotes element-wise multiplication, *W* and 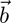 are trainable weights and biases (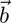 is broadcast across rows), the summation is row-wise and combines the weighted summaries from different pairs, and *f* is a stack of fully connected layers. This strategy loosely resembles a graph convolution where a grid cell is connected to every genotype-pair and edges are weighted to reflect spatial distance. Except, in our case the spatial weightings are determined by a neural network, similar to the continuous-filter convolution proposed by Schütt et al. (2017) for applications in quantum chemistry. (In practice, the number of tensors is kept to a minimum by applying equation (1) to larger tensors representing all grid cells at once: (i) a tensor of spatial scores for all cells, and (ii) a tensor built by copying *G* a total of *w*^2^ times; see Figure S2).

Last, we reshape the output to be map-shaped with separate channels for dispersal and density (resulting in a *w* × *w* × 2 tensor). Therefore, a shared set of features are used to estimate both the dispersal and population density channels of the output map. While this strategy might at first seem limiting, extracted features may flexibly align with either or both maps during training, and thus do not depend on prior assumptions about how many features each map should use. See Figure 2 and Figure S2 for additional network specifications.

The resulting model is efficient in terms of complexity: the number of parameters does not change with increasing numbers of pairs or map size. However, the computational time and memory requirements are affected by the input and map sizes. For working with large sample sizes, we offer two recommendations. First, practitioners may simply analyze a random subset of *k*_init_ pairs. Second, *k*_init_ may be further reduced to a random set of *k*_extract_ pairs, and the excluded pairs are masked from optimization of the weights in the first branch of the network. This still allows information from all *k*_init_ pairs to propagate through the network in the forward pass, a strategy that led to little or no reduction in accuracy in Smith and Kern (2023). For example, we found that using *k*_init_ = 450 and *k*_extract_ = 100 with *n* = 100 (whereas 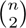 would be 4950 pairs) led to good validation accuracy in our analysis; however, we recommend exploring different numbers of pairs and testing on simulated data, as the optimal values will depend on the study system. We suggest keeping the map resolution fairly coarse, e.g., 10 ≤ *w* ≤ 50. In addition to increasing memory requirements, using a higher resolution map is not expected to improve the analysis unless spatial sampling is very dense. Furthermore, if the target maps are generated by recording density and dispersal during simulations, the map resolution may become limited by the data production step instead of RAM because it is challenging to record density with fine-resolution. Although the model quickly downsamples the genotype input using convolution and pooling, additional memory will be needed to input larger numbers of polymorphisms.

### 2.3 Training protocol

As mentioned above, we train `mapNN` on simulated datasets. Therefore, the training targets can represent either (A) the simulation-input maps conveying the carrying capacity and dispersal parameters, *K* and *σ*; or (B) maps of realized density and dispersal rate recorded during each simulation. These may be different because of demographic stochasticity and because there is not in general an exact correspondence between the parameters labeled “carrying capacity” or “dispersal” in a model and the emergent quantities in the simulation. In each of the below experiments we trained on the input maps because it is more straightforward and computationally faster than tracking realized density and dispersal during the simulation. Input maps are preprocessed before training: map values are log transformed, and the *σ* and *K* channels are independently centered and scaled. We used mean squared error (MSE) loss, therefore each grid cell in both channels of the *w* × *w* × 2 target contribute equally to the loss:

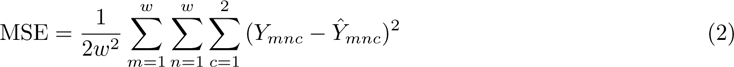

For training our neural network we used 20% of training datasets for validation, a batch size of 10, a learning rate of 10^−4^, and the Adam optimizer. We used an 80GB NVIDIA A100 GPU for training `mapNN`. Weights were initialized from a uniform distribution and biases were initialized as zero (default). The learning rate was halved whenever 10 epochs passed without reduction in validation loss, and training proceeded until 100 epochs passed without reduction in validation loss. Other settings are analysis-specific and are described below.

### 2.4 Benchmarking

#### 2.4.1 Simulation settings

Simulations used a genome size of 10^8^ bp, a recombination rate of 10^−8^ per bp per generation, and a 50 × 50 habitat. As “prior” distributions for carrying capacity and *σ* we used log-uniform on (4, 40) and (0.73, 3.08), respectively, which led to the following useful outcomes. (i) Combining even the smallest carrying capacity and dispersal rate from the proposed ranges supports local populations and avoids geographical ‘dead’ zones. (ii) The chosen ranges generated sufficient signal for the neural network to perceive and learn from. Since all areas of the map supported life (and sampling was uniform with respect to space), the neural network was not able to “cheat” by tracking local sampling density. To achieve the aforementioned range for *σ*, we fixed *σ_m_* and *σ_c_* both equal to 1, and values for *σ_f_* were calculated to obtain the desired, local *σ* according to 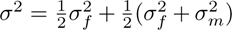. The initial population size was the mean *K* value from the input map times the squared map width. These simulations were run for 10,000 cycles in `SLiM` and recapitated in `msprime` using *N* from the final `SLiM` cycle as *N _e_*.

We chose to train on the `SLiM` input-maps because they were cleaner; the realized *σ* and density maps contained artifacts that might confuse the network. Also, it was computationally expensive to track density during the simulation, so this strategy saved much computation time. However, we did track density and dispersal throughout the simulation for 1,000 datasets used as the ground truth for evaluating our method and comparing it with existing methods.

#### 2.4.2 mapNN

We trained `mapNN` on 50,000 simulated datasets. Each dataset consisted of 5,000 SNPs from 100 individuals, with individuals sampled by uniformly selecting four individuals from each cell on a 5 × 5 grid. Training used *k*_init_ = 450 and *k*_extract_ = 100 pairs (out of 4950 possible pairs). To quantify the performance of `mapNN` we calculated the relative absolute error for each grid cell of the predicted map relative to the true map and averaged across cells to get the mean relative absolute error (MRAE) for each test dataset. We calculated separate MRAE’s for dispersal and density maps to evaluate potential disparities between the two parameters. To compare the overall performance of `mapNN` with that of `FEEMS` and `MAPS`, we took the average MRAE over 1000 test datasets. For each test, the same set of sampled individuals was analyzed with each of the different methods.

#### 2.4.3 FEEMS

We chose to include `FEEMS` in our methods comparison because it and its counterpart `EEMS` are commonly used spatial demographic inference methods that use accessible SNP input. `FEEMS` requires a user-defined triangular lattice as input, which we constructed to have 80 nodes (triangles have sides of length 6.415). The hyperparameter *λ*, which controls how constant or smooth the output will be, was set to 1. We observed better performance with `FEEMS` using a minor allele frequency cutoff of 0.05, compared to including rare alleles; this approach was used by Marcus et al. (2021) in their empirical application. The filter left between 1,556 and 2,225 out of 5,000 starting SNPs, and a similar proportion for 50,000 SNPs: between 16,083 and 22,053. We provided `FEEMS` an input table consisting of a single genotype for each individual for each SNP consisting of 0s, 1s, and 2s representing the count of the minor allele. The program was run with default settings otherwise. To calculate performance metrics for `FEEMS` we compared the estimated values along each edge in the `FEEMS` resistance grid with the true dispersal map. We averaged the dispersal rate across grid cells in the true map traversed by each edge, and used the mean dispersal rate among those cells as the ground truth for comparison with the `FEEMS` output for the edge.

#### 2.4.4 MAPS

This experiment represents a direct comparison between `mapNN` and `MAPS` (Al-Asadi et al., 2019), as `MAPS` estimates the same values as `mapNN`: dispersal rate and population density. However, `MAPS` requires accurately- defined identity by descent (IBD) blocks as input which are not available for most species, whereas `mapNN` can be used with sparse genomic marker loci. We highlighted this qualitative difference by providing `MAPS` with a relatively small number of marker loci, 5,000, and challenged `MAPS` to make use of the IBD blocks inferred from the sparse markers using empirical software. Although this experiment may seem contrived, datasets with hundreds to thousands of SNPs are much more common than IBD blocks for study systems in ecology and evolutionary biology.

The genetic input to `MAPS` are identity by descent blocks, which must be statistically inferred from empirical sequencing data. Thus, with the aim of including realistic detail in our methods comparison, we used empirical inference software to call identity by descent blocks from the SNPs of each simulated test set. For this we used the pipeline provided by the `MAPS` authors (https://github.com/halasadi/ibd_data_ pipeline), which utilizes BEAGLE (Browning and Browning, 2011, 2013), and follows the post-processing steps from Ralph and Coop (2013). `MAPS` was run using with 100 demes.

To quantify accuracy, we converted the migration rates and population sizes output by `MAPS` to dispersal rates and local densities following the recommendations from Al-Asadi et al. (2019). Specifically, the population size for a deme was divided by the deme area (habitat area / number of demes) to obtain local density, and the square root of migration rate was multiplied by the step size between demes to get *σ*. MRAE was calculated as above.

### 2.5 Empirical application: North American grey wolves

#### 2.5.1 Study system

As an empirical demonstration we applied `mapNN` to the North American grey wolf genotyping array data from Schweizer et al. (2016) with sampling localities distributed across Canada and North America. This is an ideal study system for demonstrating `mapNN` because North American grey wolves have a large range with environmental heterogeneity, and they have a moderate population size that makes spatial simulations tractable. In addition, a demographic map was estimated in this population previously by Marcus et al. (2021), allowing the dataset to serve as a benchmark for comparing inference methods. We analyzed a subset of 94 individuals characterized by Schweizer et al. (2016) as being not closely related and that were collected from 93 different sampling locations.

#### 2.5.2 Map projection

For our empirical application we generated new training data. First, we projected the habitat boundary and wolf sample coordinates from the surface of the Earth onto a planar surface for compatibility with `SLiM` simulations. To obtain an outline of the study area, we used the `map` function from the `maps` library in `R` using the Albers projection on a spheroid, while limiting the latitudinal and longitudinal ranges to (50^◦^N, 83^◦^N) and (172^◦^W, 80^◦^W), respectively. This projection preserves area between regions, but distorts shapes (regions in more extreme latitudes are stretched in the east-west dimension); however, these distortions are minimal for the current study area. The `maps` library calculated the plotted land area to be 11,083,509.65 km^2^, which was 21.93% of the complete, square map. Thus, the total map corresponds to 50,540,399.68 km^2^, with a width of 7109.18 km. The sample locations were then projected onto the map using the `mapproject` function from the `mapproj` library. We did not simulate habitat outside of Canada and Alaska; namely, parts of the population’s distribution south of the Canada-US border and in Greenland.

#### 2.5.3 Population model

To simulate wolves we used a continuous space model similar to the one described above, but with the following differences. First, we adjusted the “prior” ranges for dispersal and density to reflect the study system. For the wolf model, we aimed to simulate local *σ* between 5 and 500 km / gen, and we set *σ_f_* = *σ_m_* = *σ_c_*; therefore, dispersal, mating, and interaction distances are on the same order. To choose a prior distribution for local carrying capacity we aimed to simulate a range of population sizes that includes the currently accepted estimate from the literature. Mech and Boitani (2003) reported estimates of 52,000-60,000 wolves for Canada and 6,000-7,000 wolves for Alaska, or up to 67,000 for the combined populations of Canada and Alaska. Using the combined land area of 11,083,509.65 km^2^, we would expect to find 6.05 × 10^−3^ individuals per km^2^ on average. With this value in mind, we simulated carrying capacities between 10^−4^ and 2.46 × 10^−2^, with the maximum carrying capacity able to support 67,000 individuals if only 25% of the land area is suitable habitat.

Another simplifying assumption of our wolf model is that adult individuals do not move. Thus, the location of an individual in our model can be thought of as the centroid of their territorial range after dispersing. In addition, it is straightforward to use our method with arbitrarily more realistic SLiM models; for example, models that include adult movement, social behavior, and other biological details. Future research will be needed to explore these additional factors.

#### 2.5.4 Post hoc map correction

In the wolf model described thus far, when local dispersal rate or carrying capacity is too small, the region will not sustain a stable population. To ensure that all parts of the map support life, we applied a post hoc correction to regions that were expected to go extinct. An appropriate correction was determined by fixing carrying capacity for a uniform map, iterating over different values for dispersal rate, and noting the minimum dispersal rate that supported a stable population. This was repeated for five different values of *K*, revealing a linear relationship between log(*K*) and log(*σ*_min_) (Figure S3).

The relationship between *K* and *σ*_min_ was used to update each training map as follows. For each map we randomly chose which parameter to boost, either *σ* or *K* ; this strategy allowed us explore maps with small local *σ* (but relatively large *K* after providing a boost) in addition to maps with small local *K* (but relatively large *σ*). For each grid cell in a given map, if the cell’s parameter combination fell below the empirically derived threshold, we applied the correction: 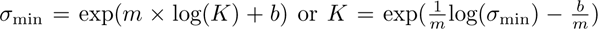, where *m* and *b* are from the linear fit of log(*σ*_min_) to log(*K*). Last, we further increased the adjusted values by 25% to allow for random noise.

While it might seem useful to try to identify uninhabitable regions, such an inference comes with additional challenges. For populations with local extinctions, sampling could not follow the grid approach or empirical sampling strategy described above, but must be opportunistic. With opportunistic sampling the network would learn to rely on sampling density rather than genetic information for inferring population density (which may be a valid source of information, but is not the goal of this study).

#### 2.5.5 Genomic resources

In our training simulations we included 38 independent genomic segments with lengths corresponding to those of dog chromosomes from the Ensembl genome database (Cunningham et al., 2022). We applied chromosome-average recombination rates calculated from the Campbell et al. (2016) dog genetic map. All sites were assumed neutral. SNPs from Schweizer et al. (2016) with missing data (and indels or non-biallelic SNPs, if present) were filtered to obtain 10,627 SNPs represented in all individuals.

#### 2.5.6 Demographic history

We simulated North American grey wolf demographic history using the two-epoch model from Fan et al. (2016) and Robinson et al. (2019), with an ancestral *N _e_* of 45,000 and North American bottleneck of *N _e_* = 17, 350. The bottleneck occurred 1,250 years before present, or 417 generations using a generation time of three years (Bergström et al., 2022). To keep simulations tractable we simulated only 1,000 years in spatial SLiM for the grey wolf analysis. In effect, our complete model has three time epochs, with the most recent generations simulated in continuous space with spatially heterogeneous demography.

#### 2.5.7 Habitat mask

Starting with the segmented training maps described above, we applied a mask in the shape of the simulated habitat. Grid cells outside of the habitat mask—over water, and south of the US-Canada border—were changed to a value of zero for both the dispersal and density channels, and the resulting maps were used as input to `SLiM` for simulating training datasets. The loss function for `mapNN` was modified to exclude cells beyond the habitat boundary. Likewise, to evaluate `mapNN` the MRAE included only cells within the habitat boundary. When plotting estimated wolf maps, values outside of the habitat were visualized as white. See Figure S4 for a sample of training maps from the empirical application.

#### 2.5.8 Sampling

For each empirical sampling location, we sampled a random individual within a radius of 2% of the map width. If no individuals were available within the starting radius, the sampling radius was doubled. This procedure was repeated until the empirical sample size was achieved. The outcome is a sampling scheme that roughly mirrors that of the empirical sample localities.

## 3 Results

### 3.1 Simulation benchmark

We evaluated the performance of `mapNN` on held-out, simulated datasets, each consisting of 5,000 SNPs from 100 individuals, with individuals sampled uniformly from a 5 × 5 grid. In each simulation, carrying capacity and dispersal rate varied across the landscape at a coarse scale. For training, we used the carrying capacity and dispersal rate simulation-input maps as targets. However, for testing we tracked density and dispersal during the simulation, and compared these ground truth maps against predictions from `mapNN` and the other methods.

Visual inspection revealed that in most cases `mapNN`’s output resembled the true maps (Figure 3). After taking the MRAE across grid cells in a map, the average MRAE among tests in this experiment was 0.17 and 0.34 for the dispersal and density maps, respectively. This means that values in the predicted dispersal map were around 17% off from the corresponding location in the true map and values in the density map were around 34% off, on average. MRAE was roughly 50% higher when either the dispersal or density map was heterogeneous, and MRAE was doubled when both maps were heterogeneous. See Figures S5-S9 for a sample of predicted maps from this experiment.

**Figure 3:**
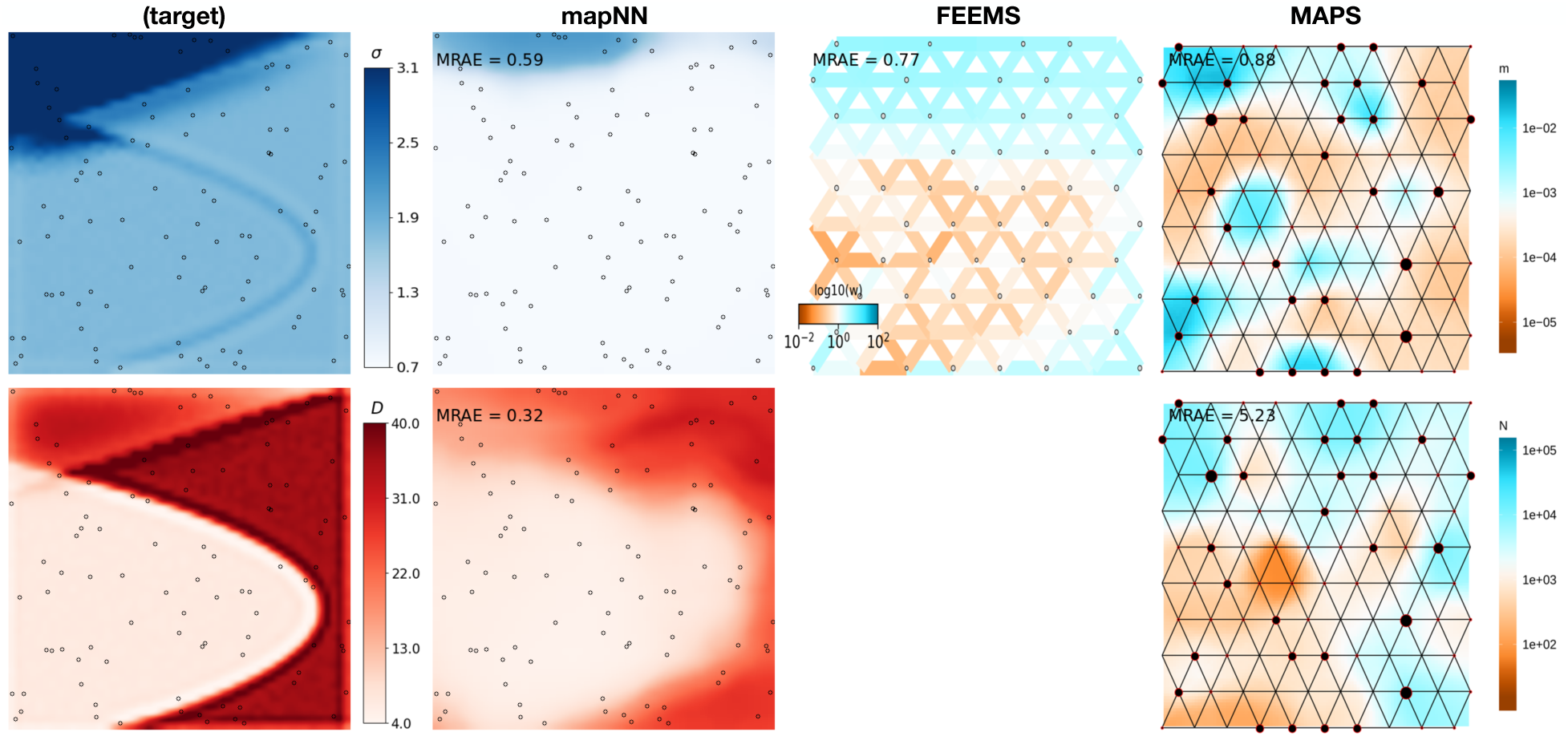
Predicted maps for an example test dataset. The leftmost column shows the ground truth maps for dispersal (top row) and density (bottom row). Columns 2-4 show estimated maps using three different methods with 5,000 single nucleotide polymorphisms: `mapNN`, `FEEMS`, and `MAPS` (respectively). Note that `FEEMS` and `MAPS` use a different color scale than mapNN and the input maps. Output from `mapNN` is smoothed (after calculating error) by resizing the map to 500×500 using the `PIL` library with cubic spline interpolation.

Next we compared our method with two existing methods for estimating demographic maps. `FEEMS` (Marcus et al., 2021) is not designed for estimating separate maps for dispersal and density, but rather to estimate a single map of effective migration which includes components of both dispersal and population density. In addition, the effective migration rates estimated by `FEEMS` are *relative* values. To directly compare our method with `FEEMS`, we looked at the correlation (instead of error) between the `FEEMS` output and the true dispersal rate map across test datasets. The performance of `FEEMS` evaluated this way was comparable with that of `mapNN`: mean *r* ^2^ between the true and inferred map values was 0.216 with `FEEMS`, 0.221 with `mapNN` for dispersal, and 0.240 with `mapNN` for density (Figure 4). As an alternative metric, we multiplied the relative values output by `FEEMS` by the range-wide mean dispersal rate from the true map to get an estimate for dispersal rate for each edge in the `FEEMS` resistance network. Using these values, the average “dispersal” MRAE was 0.26.

**Figure 4:**
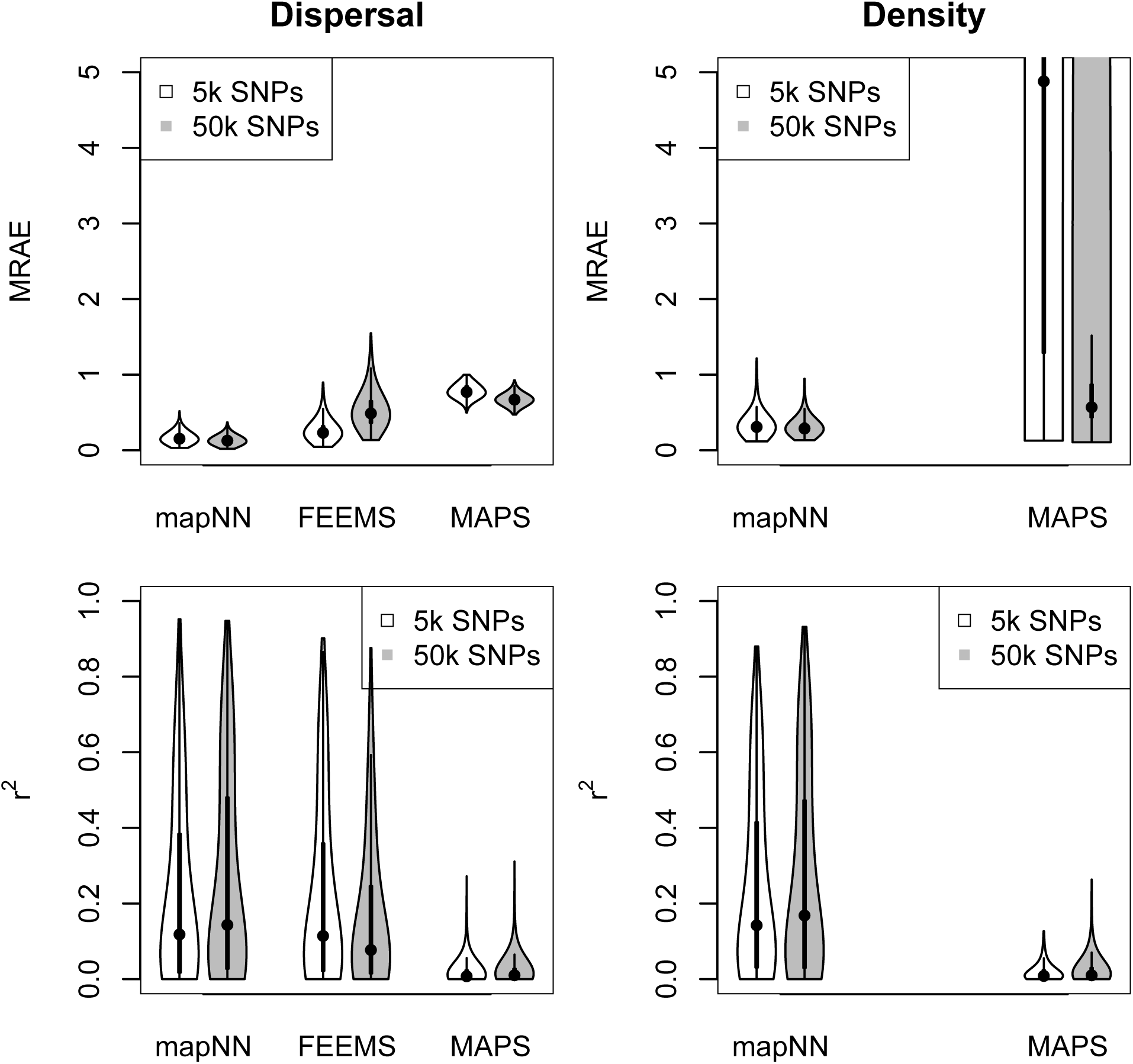
The top row shows the map-wide mean relative absolute error (MRAE) across 1000 test simulations. The density plot is truncated for visualization. The violin plot for `MAPS` with 50k SNPs has a large proportion (22%) of values above the 75th quantile + 1.5 × the interquartile range (commonly used to define outliers). `FEEMS` errors are shown in the “dispersal” panel for convenience, although `FEEMS` technically estimates effective migration. The bottom row shows the correlation (*r* ^2^) between the estimated and true maps.

The `MAPS` program (Al-Asadi et al., 2019) was not expected to fare well in this experiment, as we provided it with inadequate genotype data for calling identity-by-descent tracts: only 5,000 SNPs. Nonetheless, the dispersal and density surfaces estimated by `MAPS` looked quite good on some of the more heterogeneous maps, where it sometimes recapitulated the spatial breaks between high- and low-density segments of the habitat. However, the magnitude of estimated density was far from the true density in most cases which led to high error: average MRAE of 0.78 for dispersal and 12.75 for density. The `MAPS` model did not converge for two test datasets, which were not included in the average MRAE.

In a separate experiment we increased the number of SNPs to 50,000, and because of memory constraints were limited to analyzing 350 sample-pairs with `mapNN` (down from 450 pairs). Using the larger genotype input incrementally improved the ability of `mapNN` to infer dispersal and density maps (average MRAE=0.13 and 0.31 across the test set, respectively). Oddly, using 50,000 SNPs with `FEEMS` caused error to increase significantly (average MRAE=0.53). We speculate that unaccounted-for linkage between adjacent SNPs may be responsible for this behavior using `FEEMS`. One dataset did not converge using the `FEEMS` algorithm and was not included. The larger input did allow `MAPS` to infer more accurate dispersal maps (average MRAE=0.67), and substantially better density maps (average MRAE=1.58; see Figure 4). However, mapNN with only 5,000 SNPs still did consistently better than `MAPS` even given 50,000 SNPs.

### 3.2 Model misspecification

The above benchmarking was done in a well specified setting– i.e., the test data were produced using the same simulation procedure and parameter ranges as the training data. That test therefore represented a best case scenario for our method. Next, we challenged the ability of `mapNN` to generalize by exploring scenarios where the test data was generated using parameter values outside of the ranges used during the training procedure, a type of model misspecification.

We found that `mapNN` predictions were mostly constrained to the range of values used in training (Figure 5). The performance of `mapNN` was poor on extreme test cases where the examined parameter values fell outside of the training distribution (i.e. extrapolation). When test datasets used a very small dispersal rate—2.70 times smaller than the minimum of the training distribution—and flat maps for both density and dispersal, estimates for dispersal rate fell near the minimum of the training distribution (red points in Figure 5). In the same experiment, `mapNN` simultaneously underestimated population density, although this was not misspecified.

**Figure 5:**
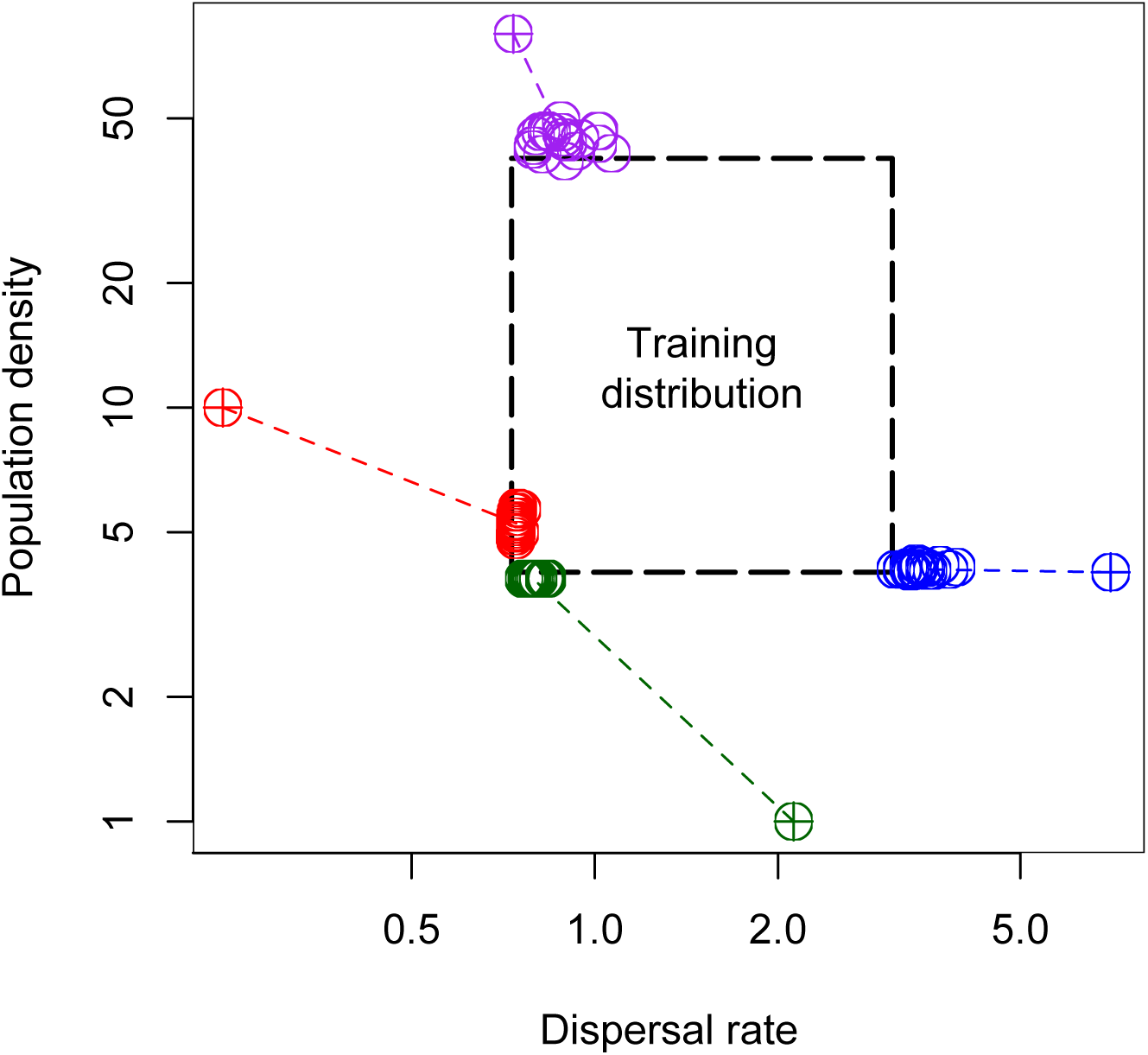
Output from four different scenarios with out-of-sample test data. The dashed, center box shows the ranges for population density and dispersal rate used during training. Colored, dashed lines connect test values (crosshairs) to predictions (open circles). Test maps in these experiments had uniform density and dispersal.

Similarly, we explored a test scenario with very large dispersal rate (1.89 times larger than the maximum from training; blue points in Figure 5). In this test `mapNN` estimated smaller than expected dispersal rate, while the density estimate remained accurate. In this experiment, `mapNN` was able to extrapolate slightly beyond the maximum from training, but was mostly constrained to the prior range.

Next, we explored scenarios with misspecified population density. The estimates from `mapNN` showed a similar pattern as before: when the test data had very small density (4 times smaller than the minimum of the training distribution), the estimates from `mapNN` fell near the minimum value from training, and dispersal rate was underestimated (purple points in Figure 5). When the test data had very large K (twice the maximum from training), the estimated density fell near the maximum from the training distribution, and dispersal rate was moderately overestimated (green points in Figure 5).

To summarize, `mapNN` is not expected to extrapolate well, and adequate care must be taken when designing and performing simulations to train a model. A potential indicator that adjustments to the training distribution are needed might be if either population density or dispersal estimates fall near or outside the training distribution. Importantly, the above results with out-of-sample test data may be specific to the examined scenario (e.g., square map, random sampling, etc.), prior ranges, and other settings (e.g., *n* = 100, 5000 SNPs, etc.).

### 3.3 Empirical application: North American grey wolves

Training simulations used a habitat shaped like North America and prior distributions that are reasonable for wolves, but were otherwise similar to the aforementioned simulation model. We attempted to run 5,000 simulations, however we limited the runs to a one week wall time. As a result, the wolf training set consisted of 2,938 completed simulations and the resulting training distribution was skewed towards simulations with smaller dispersal rate and smaller density; the largest completed simulations had a total population size exceeding 140,000, which is roughly double the current population size estimate (Mech and Boitani, 2003). Next, we augmented the training set by taking ten sets of sampled individuals from each simulation with replacement. The resulting training set consisted of 29,380 datasets. For training we included a subset of genotype pairs (*k*_init_ = 650 and *k*_extract_ = 100), and masked grid cells falling outside of the habitat from the loss function (see Methods).

Validation on simulated data gave MRAE=0.26 for the dispersal map and MRAE=0.44 for the population density map. These errors were higher than the above benchmark analysis, which may be attributable to using larger “prior” ranges for the unknown variables, uneven sampling across space, a 40% smaller training set, and 10x fewer unique training maps. See Figures S10-S14 for a sample of validation plots using simulations from the grey wolf analysis.

For the final, empirical prediction we estimated 100 maps using different, random subsets of genotype pairs and used the average across replicates as a point estimate for each cell in the map grid (Figure 6A). The resulting demographic map had a range-wide mean dispersal rate of 68.8 km per generation. The inferred dispersal map also varied across space between 58.0 km per generation in southern latitudes to 242.6 km per generation in Northern Canada. The range-wide mean density from the estimated map was 11.6 individuals per 1000 km^2^, which gives a total population size of 129,302 for the studied region; however this value is likely inflated by highly uncertain density estimates in Eastern Canada where sampling was sparse. Estimated density ranged from one individual per 1000 km^2^ in Northern Canada and Western Alaska to 100 individuals per 1000 km^2^ in Eastern Canada.

**Figure 6:**
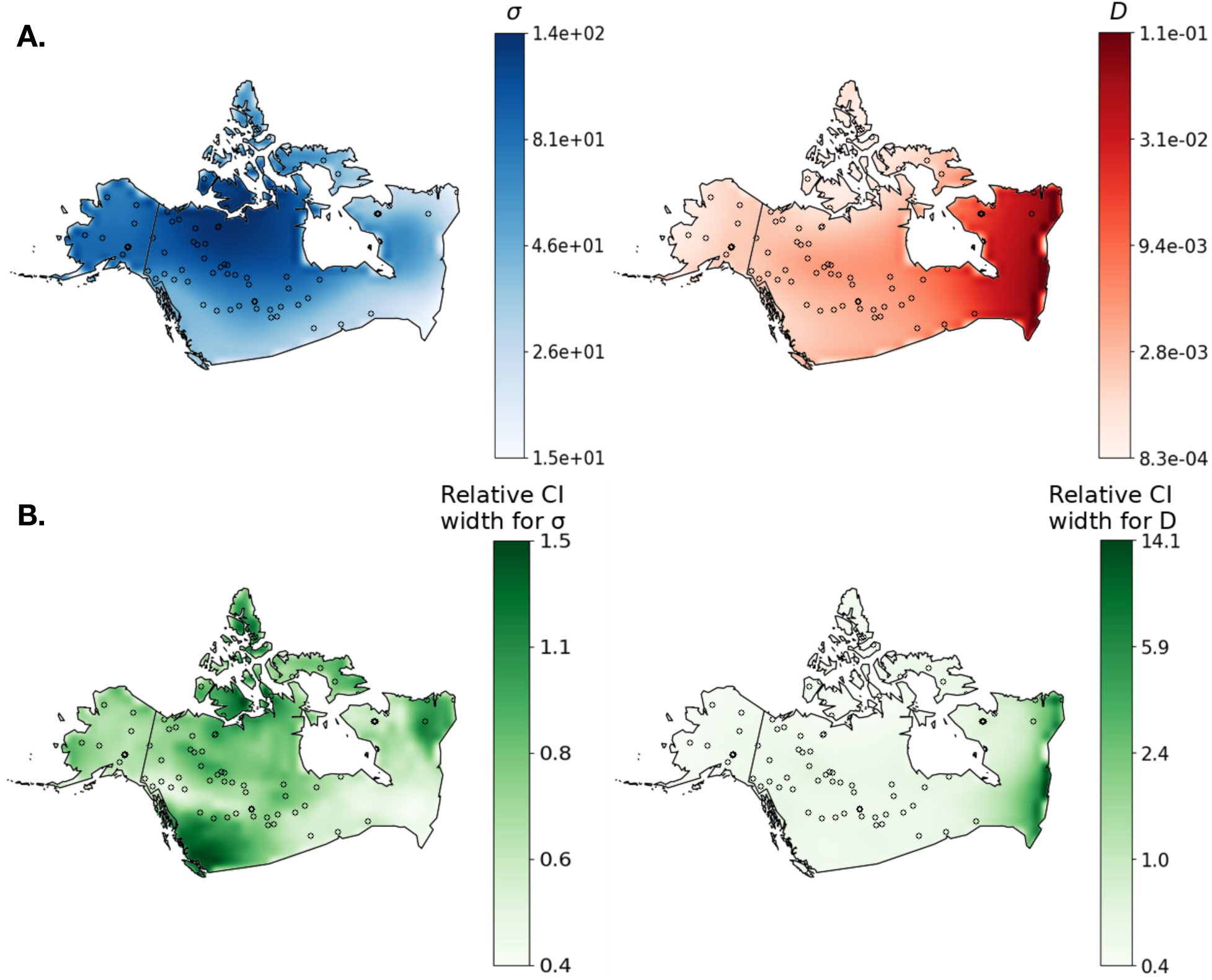
(A) Maps estimated from grey wolf data from Schweizer et al. (2016). The left panel shows estimated dispersal rate (km / generation) and the right panel shows estimated density (individuals / km^2^). (B) Uncertainty maps: each heat-map conveys the width of the 95% confidence interval for each grid cell from parametric bootstrapping. Note the color bar ticks have log spacing.

To quantify uncertainty we performed a parametric bootstrap procedure by simulating 100 additional datasets under the inferred dispersal and density maps, and re-estimating demography for each replicate. Treating each demographic parameter—dispersal rate and density—separately, we calculated 95% confidence intervals for each grid cell in the map as *t*^∗^_0.975_ − *t*^∗^_0.025_, where *t*^∗^*_α_* is the *α*th quantile of the resampled distribution for the cell (Davison and Hinkley, 1997). Figure 6B shows the width of the interval for each grid cell relative to the point estimate *t*, (*t*^∗^_0.975_ − *t*^∗^_0.025_)*/t*, across the studied region, which hence summarizes (relative) confidence. Relative uncertainty of dispersal rate was highest in British Columbia, and lowest in parts of Quebec. Meanwhile, confidence in population density was relatively uniform across space, except for Eastern Canada where there was moderate (in Ontario) to very high (on the Atlantic coast) uncertainty. Coinciding with high uncertainty, there was also a lack of sampling locations in Southeast Canada. Across the whole habitat, unsampled areas of the habitat had on average 37% wider relative confidence intervals for population density than areas with at least one sample (using non-overlapping 5 × 5 grid cell windows). The same was not true for dispersal rate confidence intervals (unsampled windows had 12% narrower intervals). We found this interesting, as it suggests quite different strategies by the neural network for estimating each parameter.

## 4 Discussion

### 4.1 Population genetics for spatial demographic inference

A longstanding aim of molecular ecology has been to estimate local dispersal rates and population densities across the landscape using population genetic data. However, currently available methods require inputs such as inferred identity by descent tracts that may not be obtainable in important non-model systems (Ringbauer et al., 2017; Al-Asadi et al., 2019). Our deep learning method for estimating dispersal and density maps is designed to use small-to-intermediate numbers of polymorphisms (e.g., 5,000) and sampled individuals (e.g., 100). Our simulation-based inference procedure also offers flexibility, as training datasets can be tailored to the empirical study system. However, the production of training datasets can also be computationally expensive and must be carefully designed to match the study system; exactly how carefully, i.e., which aspects of demography are most important to match, is currently unknown. Therefore, we expect our method to be particularly useful for smaller populations with incomplete genomic resources, indeed as is often the case for conservation targets.

In our benchmark analysis on simulated data, `mapNN` outperformed existing methods `FEEMS` and `MAPS`, making `mapNN` a state-of-the-art tool for spatial demographic inference. However, this comparison was tricky due to qualitative differences between the examined programs’ input types and the quantities they estimate. In particular, the `MAPS` program (Al-Asadi et al., 2019) is intended for high depth, high marker density datasets, such that accurate identity-by-descent blocks may be statistically inferred; estimating such blocks is a preliminary step in the `MAPS` pipeline and is itself error prone. Thus, `MAPS` was not expected to fare well with the moderate number of polymorphisms we supplied it with (up to 50k), and our analysis indeed showed that `mapNN` is more accurate with relatively sparse genotyping data. Nonetheless, it was encouraging that `MAPS` was still able to detect spatial heterogeneity in some tests, even with the smaller numbers of SNPs used as input. The ability of `mapNN` to work with reduced representation or low-coverage whole genome datasets makes it a valuable resource for non-model species with limited data.

The comparison with `FEEMS` (Marcus et al., 2021) was less direct, as the values output by `FEEMS` are relative migration rates instead of the dispersal rate and density parameters that `mapNN` estimates. In effect, `mapNN` and `MAPS` have a more challenging task because they estimate the *magnitude* of dispersal and an additional map for the separate density parameter. Even so, in our simulation study `mapNN` had lower error than `FEEMS`. We acknowledge, however, that different test scenarios than the one we used may produce a different outcome. The error values reported in our analysis are specific to the test data we chose to analyze, and will depend on the settings and specifications of each respective method. In summary, `FEEMS` remains a very attractive tool for situations where only relative migration rates are sought for evaluating landscape connectivity because it doesn’t require simulating training datasets. However, to obtain actual values for dispersal rate and density from genetic data, methods like ours and `MAPS` are more appropriate. Also, our method can more easily take into account different aspects of biological realism, for instance different rates for parent-offspring dispersal and mating dispersal, directionally biased dispersal, or temporally varying dispersal.

One drawback of most inferences in spatial population genetics is ecological interpretability. For example, the methods of Rousset (1997), Ringbauer et al. (2017), and Al-Asadi et al. (2019) estimate effective dispersal, which is influenced by both natal dispersal and the distance between mates. Similarly, the inference of effective population density rarely translates to the census number of individuals per unit area. As a result, the effective values inferred by conventional population genetic methods do not have a clear ecological interpretation. Excitingly, simulation-based inference may provide a means to estimate *ecological* parameters using genetic data if enough information is known about the study system. For example, if the mother-offspring distance is known—e.g., the distance that seeds fall from the plant is predictable in some species—then the mating distance—e.g., pollination—could potentially be directly inferred using population genetic data. Likewise, if the simulated fecundity and other relevant variables match that of the studied population, our inference methods may move beyond *N _e_* to estimate actual population density. Exploring these possibilities will require additional research.

### 4.2 Alternative approaches for analyzing spatial demography

Whereas `mapNN` is designed to estimate effective population density (and dispersal) some generations in the past, the actively developing research area of close kin mark recapture (CKMR) may be used to directly estimate the size of a population (Bravington et al., 2016). CKMR generally involves genotyping to identify parent-offspring or sibling pairs in a single temporal sample, and the ratio of close kin to total samples can provide information about population size, having made assumptions about the family size distribution of the population. However, an important assumption of most current CKMR analyses is that all individuals of a particular age have an equal probability of being sampled (Conn et al., 2020), which in practice may to lead to biased population size estimates, particularly for widely distributed species with limited dispersal. CKMR generally requires large sample sizes, with sample size scaling with the square root of abundance (Bravington et al., 2016), however much larger sample sizes would be required to analyze wide ranging species, e.g., North American grey wolves.

Population genetics-based approaches like ours can use relatively small sample sizes. Furthermore, as genomic sequencing data continue to be produced for separate molecular ecology applications, for example, studying population structure, the same data may be reused for spatial demographic inference using `mapNN`. Moving forward, our approach may be used with polymorphisms from eDNA and non-invasive sampling (Adams et al., 2019; Andres et al., 2023). In addition, simulation-based inference allows researchers to tailor training simulations to their respective study system. For example, training simulations for `mapNN` can be customized to mirror the irregular habitat shapes we find in nature; this challenging variable will otherwise cause biased estimates. Another advantage of population genetics based approaches in this context is estimating connectivity and dispersal surfaces across space, for which there are limited tools using close kin data; of course, this comes with the downside that it is challenging to estimate present day demographics using polymorphisms.

### 4.3 North American grey wolf population biology

We estimated coarse-grained maps of population density and dispersal in North American grey wolves using data from Schweizer et al. (2016), and found that our estimated maps were in rough agreement with previous research. Our estimated maps showed spatially heterogeneous dispersal, with relatively low dispersal rates in British Columbia, across Southern Canada, and in Northern Quebec; and several times higher dispersal in north-central Canada. This outcome is consistent with the observation by Mech and Boitani (2003) that territory size correlates positively with latitude; for example, the smallest territory in the studied region was 149 km^2^ in Algonquin Park in South Ontario, and the largest was 1,645 km^2^ in South-central Alaska Mech and Boitani (2003). The range-wide mean dispersal rate estimated by our analysis, 68.8 km, was noticeably lower than the expected dispersal rate between territories of 98–147 km/generation measured using radio collars (Jimenez et al., 2017; Kojola et al., 2006; Barry et al., 2020). However, this outcome was expected, because the mean per generation dispersal rate that we estimate is a function not only of between-territory dispersal, but also of non-dispersing wolves, the reproductive success of dispersing wolves, and other factors, and should be smaller than the between-territory average.

The inferred map showed lowest density in Alaska and the far north of Canada, gradually increasing density through Central Canada, and very high population density in Eastern Canada. We compared our estimated density map with data summarized by Mech and Boitani (2003) showing approximate wolf numbers by Canadian province and U.S. state. It is important to note that the wolf numbers reported by individual province are approximate census estimates, and they vary on a smaller scale than the coarsely-segmented maps we used for training. Considering the northerly regions together, Alaska and the Canadian provinces of Yukon, the Northwest Territories, and Nunavut combined had the lowest reported density from Mech and Boitani (2003), 4.4 individuals / 1000 km^2^ (although Yukon had the highest reported density of any province, 10.5 individuals / 1000 km^2^). Consistent with these previously reported numbers, the lowest densities in our estimated map were in Alaska and Northern Canada, and ranged between 1 and 7 individuals / 1000 km^2^, approximately. Focusing on another set of provinces in South-Western and South-Central Canada—British Columbia, Alberta, Saskatchewan, and Manitoba—the reported density from Mech and Boitani (2003) was 7.9 individuals / 1000 km^2^. Our estimated map was again reasonably accurate, where for this region our estimates were roughly 3-7 individuals / 1000 km^2^. However, in the remaining eastern provinces of Ontario, Quebec, and Labrador (excluding Newfoundland), the reported density from Mech and Boitani (2003) was 5.9 individuals / 1000 km^2^. The latter region was markedly different in our estimated map, which showed very high density in Eastern Canada. But, uncertainty was also very high in Eastern Canada, likely due to a lack of sampling in that region. Our total population size estimate of 129,302 was larger than the census estimate of 58,000-67,000 (Mech and Boitani, 2003); again, the total size was inflated by the uncertain density estimates in Eastern Canada.

In summary, we expect the demographic maps estimated by our program to be reasonably accurate, except for under-sampled regions where population density was highly uncertain. There are several reasons for the dissimilarities between our estimates and the reported values from Mech and Boitani (2003). Besides the issue of non-uniform sampling, we also ignored the social dynamics of wolves, and assumed that demography was constant throughout the recent past. We further speculate that the high density point-estimates in Quebec and Ontario might reflect gene flow from nearby populations in the United States in Minnesota and around the Great Lakes. Repeating the analysis with more fine-grained spatial segments might allow for a more direct comparison with the Mech and Boitani (2003) data.

Marcus et al. (2021) also analyzed the North American grey wolf data from Schweizer et al. (2016), estimating effective migration surfaces using various levels of spatial smoothing (by tuning `FEEMS`’s *λ* parameter). The smoothest map estimated by Marcus et al. (2021) (*λ* = 100) is perhaps most comparable to the coarsely-segmented training maps we used with `mapNN`. At this level of smoothing, the `FEEMS` map showed a subtle reduction in effective migration through South-western and South-central Canada. Thus, inferred (coarse) maps from `mapNN` and `FEEMS` are weakly correlated but there are notable differences. For instance, Eastern Canada had lower-than-average dispersal with `mapNN` but higher-than-average effective migration with `FEEMS`. To reiterate, the values estimated by `FEEMS` are impacted by both dispersal and density, whereas our method aims to disentangle the two values. The other `FEEMS` plots inferred by Marcus et al. (2021) used finer spatial granularity, and, if they are more accurate than the smoother map, could offer an alternative perspective on effective migration in this population. However, it is unclear if the fine scaled migration barriers identified by `FEEMS` are real or analysis artifacts, as the sampling was not very dense, and `FEEMS` is not free of error (Marcus et al., 2021; Lundgren and Ralph, 2019). This comparison has another complication due to a difference in sample size: the Marcus et al. (2021) analysis used additional specimens (*n* = 111). To facilitate a direct comparison between the two programs, `FEEMS` was re-run on the *n* = 94 dataset used in the current study by our colleague, Vivaswat Shastry (Figure S15; personal communication). The new map (with *λ* = 100) was qualitatively similar to the original FEEMS run, supporting that the difference between outputs from `FEEMS` and `mapNN` is due to divergent modeling approaches rather than sampling effects.

An additional shortcoming in our grey wolf application was that we ignored parts of the species distribution. In particular, significant grey wolf populations exist in Minnesota and other areas below the Canada-U.S. border, and approximately half of the species exists in Eurasia (Mech and Boitani, 2003) with the potential to contribute migrants to North America. This misspecification may bias demographic estimates using our approach, as shown by Smith et al. (2023) for estimating dispersal rate. We judged this to be a reasonable starting point because populations south of the Canada-U.S. border are smaller than the analyzed populations, and gene flow with Eurasia is very limited (Sinding et al., 2018). However, we did estimate a larger than expected population size, which could be related to the additional grey wolf populations.

### 4.4 Additional limitations

The quality of predictions from `mapNN` depends strongly on details of the training data, in particular whether they are similar to the test (empirical) data. Above we discussed how ignoring parts of the species range in the training simulations can bias predictions. In addition, we explored the ability of `mapNN` to extrapolate beyond the training distribution by testing on datasets with extreme population density or dispersal rates. We found that predictions were mostly constrained to the range of parameters covered by the training distribution, although some estimates fell just outside the range of values used in training. This suggests that shortcuts are not available for these parameters: it is not sufficient to simulate a down-scaled or local population as training data. Therefore, particularly for populations with very large present day size, it is currently challenging to simulate appropriate training datasets. One potential strategy to address this is to simulate fewer generations that are spatially explicit, as we have done in the current study and in Smith et al. (2023), which approximates initial genetic diversity as having been produced by random mating. This strategy may work for some situations better than others, and researchers should use simulations to validate the approach. Using this shortcut, a subset of our simulations ran to completion with more than 100,000 individuals. However, population size is not the only variable affecting simulation runtime. In particular, as the mating and competition distances increase, the number of interactions increases exponentially and the simulation is slowed further. As an approximate guideline, it should be possible to simulate populations under 100,000 individuals with moderate interaction distances. Although, much larger populations can be simulated if the interaction distance is small, and the runtime will ultimately depend on the simulation model.

Details of the training map will also influence the performance of `mapNN`. Our current training maps had relatively few spatial segments representing different environmental conditions, and the resulting neural network learned to predict maps with similarly coarse segmentation. We expect that using training maps with more, smaller segments would allow the model to detect smaller spatial details—up to a point, at least. Of course, what resolution can be inferred will depend strongly on sampling density and other analysis parameters. A separate idea is to use data from the real world to inform the training maps. For example, this could involve using GIS layers of environmental variables to guide the scale of spatial heterogeneity, or eventually incorporating environmental data into the analysis. Along such lines, the integrative distributional, demographic and coalescent (iDDC) approach combines distribution modeling via environmental inputs with coalescent simulations and can be used to analyze spatially heterogeneous density (Lacey Knowles and Alvarado-Serrano, 2010; Brown and Knowles, 2012; He et al., 2013).

There is generally not enough signal in polymorphism data to detect very recent population size changes; although the temporal resolution of demographic inference will depend on the population size, mutation rate, number of variants, and other variables. Therefore, models that assume a constant *N _e_* actually estimate some form of average, historical *N _e_*, and not necessarily the present day *N _e_*. Multiple-time epoch models may do a better job at estimating recent *N _e_*, but are still constrained by a lack of information about recent times. Our analysis likely suffers from this shortcoming as well, as our training data assumed constant demography for the most recent 1,000 years and we use a relatively small number of SNPs. It is promising that more recent population sizes can be inferred using linkage and identity by descent information (Ringbauer et al., 2017; Al-Asadi et al., 2019; Santiago et al., 2020; Browning and Browning, 2015; Palamara et al., 2012) (MSMC), and the field is drawing continually closer to estimating contemporary population sizes from genetic data.

Another potential source of model misspecification is sampling strategy. Smith et al. (2023) showed that if the training data use uniform sampling, for example, but the test data use asymmetric or other non-uniform sampling schemes, then predictions will be biased. However, if the training simulations were sampled intentionally to match the sampling scheme of the test data, then accuracy was restored. A separate issue may arise when analyzing heterogeneous density: if sampling chooses individuals uniformly from the population, more dense regions get sampled more heavily, and so the neural network would undoubtedly learn to predict local density directly from sampling density, which could inhibit the learning of relationships between genotypes and density. The above issues can by circumvented by following the empirical sampling scheme we used when designing training data.

Although we expect simulations to be the most computationally expensive step, we also ran into challenges related to training the `mapNN` deep learning model. All but the smallest analyses will require a GPU, as our training was prohibitively slow using as many as 50 CPU cores. Moreover, memory may be limited on a GPU: we used a subset of pairs, as not all pairs of individuals fit into memory. The default behavior of the neural network involves looping over all combinatorial pairs of genotypes, the number of which increases quadradically as the sample size increases. For reference, we analyzed a subset of 350 or 450 (for 50,000 and 5,000 SNPs, respectively) pairs out of 4950, on an 80 Gb GPU.

### 4.5 Summary

We present a polymorphism-based method for estimating spatial maps of population density and dispersal rate. This addresses a gap in available methods, as similar tools either require identity-by-descent blocks or estimate relative migration surfaces. As with many other machine learning models, `mapNN` was challenged to extrapolate beyond the training distribution; therefore, workflows will require custom simulations for analyzing each new population or study system. Our analysis in North American grey wolves produced reasonable outputs, but accuracy was affected by spatial sampling density. We expect `mapNN` to be most useful for small to moderate sized populations that are feasible to simulate, for example, many endangered species.

## Acknowledgements

We thank Vivaswat Shastry for the supplementary `FEEMS` run on the North American grey wolf data, John Novembre for important feedback on the descriptions of `EEMS` and `FEEMS`, and members of the Kern-Ralph colab for feedback throughout the project. This work was supported by the National Institutes of Health awards F32GM146484 to C.S. and R01HG010774 and R35GM148253 to A.D.K.

## Data Accessibility and Benefit-Sharing Section

The `mapNN` code is available at https://github.com/kr-colab/mapNN. Benefits from this research arise from the sharing of our software, as described above. Data for North American grey wolves were published by Schweizer et al. (2016) and are available from https://datadryad.org/stash/dataset/doi:10.5061/dryad.c9b25.

## Supplementary material

**Figure S1:**
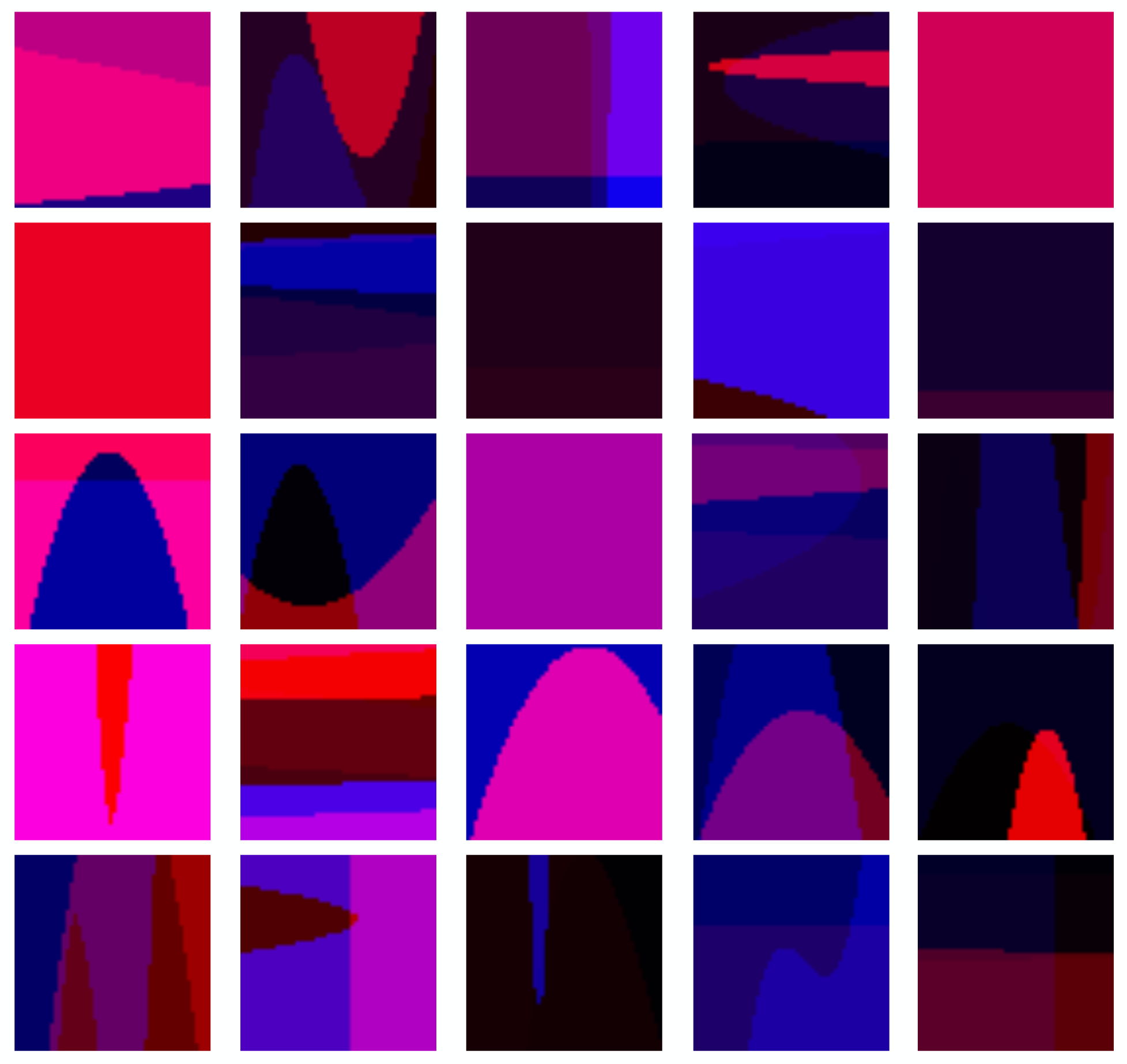
PNG renderings for a random selection of training maps for the benchmark analysis. The blue channel conveys dispersal rate and the red channel conveys carrying capacity.

**Figure S2:**
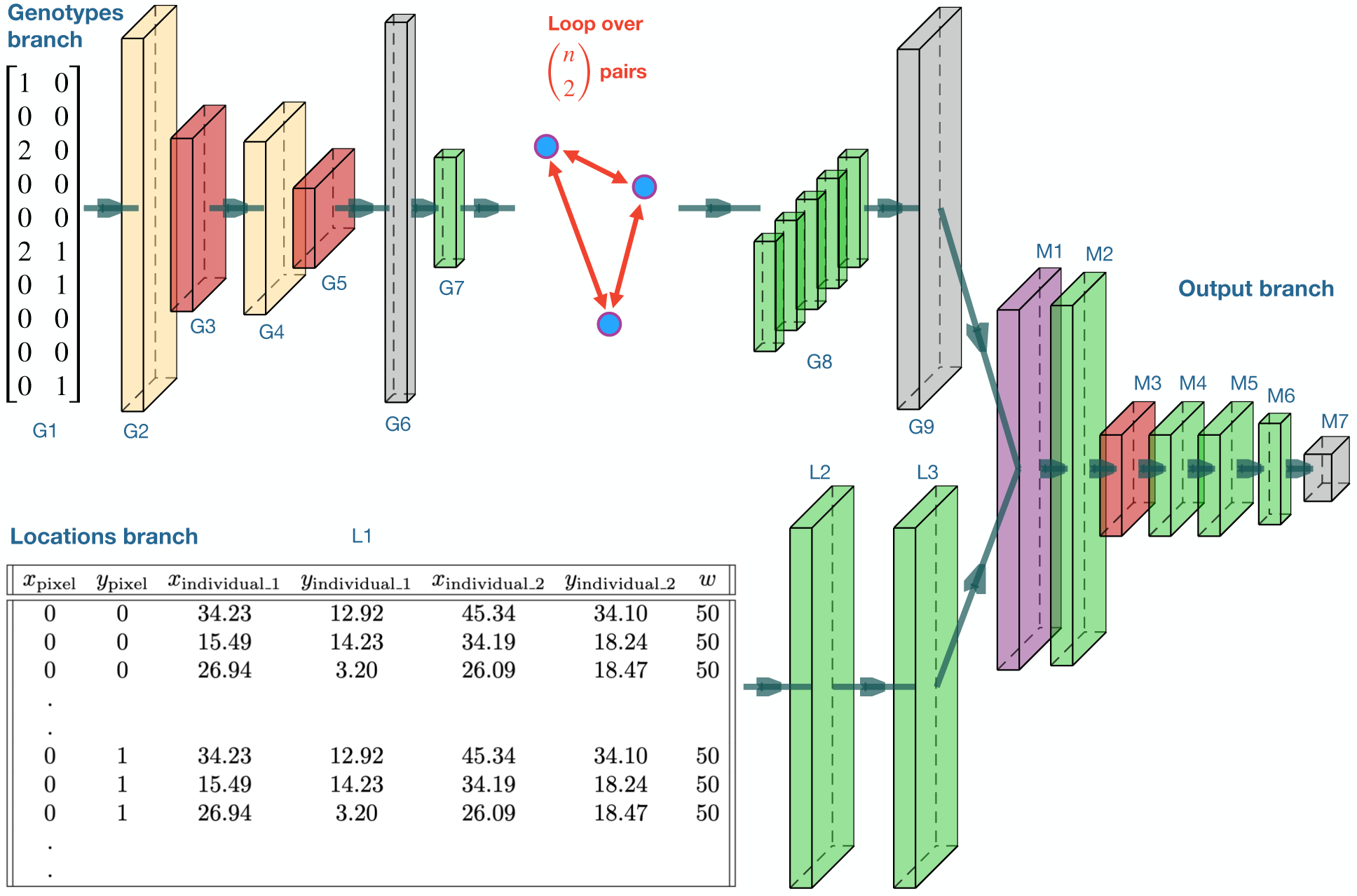
Same diagram as Figure 1, with extended caption for descriptions and output sizes of each layer. Visualized tensor sizes are proportional to the cube root of actual dimensions if *s* = 5, 000 SNPs, *k* = 10 pairs, and *w* = 10 map width were used. • G1. (*s*, 2) Genotypes for a pair of individuals. This branch of the network will be repeated for multiple pairs. • G2. (*s*, 64) 1D convolution, kernel size 2, 64 filters, rectified linear unit (ReLU) activation. • G3. (*s/*10, 64) 1D average pooling, window size 10. • G4. (*s/*10, 108) 1D convolution, kernel size 2, 64 filters, ReLU. • G5. (*s/*100, 108) 1D average pooling, window size 10. • G6. (108*s/*100) Flatten. • G7. (128) Dense, 128 filters, ReLU. • G8. (128, *k*) Outputs from looping over pairs (only five pairs are shown). • G9. (*kw*^2^, 128) The outputs from *k* pairs are stacked together, and then duplicated for each of *w*^2^ grid cells. • L1. (*kw*^2^, 7) Locations table for every combination of grid cell and genotype-pair (not all rows are shown). • L2. (*kw*^2^, 128) Dense, 128 filters, ReLU. • L3. (*kw*^2^, 128) Dense, 128 filters, ReLU. • M1. (*kw*^2^, 128) Element wise multiplication between layers G9 and L3, followed by ReLU. • M2. (*kw*^2^, 64) Dense, 64 filters, ReLU. • M3. (*w*^2^, 64) 1D pooling on every *k* rows. • M4. (*w*^2^, 64) Dense, 64 filters, ReLU. • M5. (*w*^2^, 64) Dense, 64 filters, ReLU. • M6. (*w*^2^, 2) Dense, 2 filters (linear activation). • M7. (*w*, *w*, 2) Rearrange into a stack of two maps.

**Figure S3:**
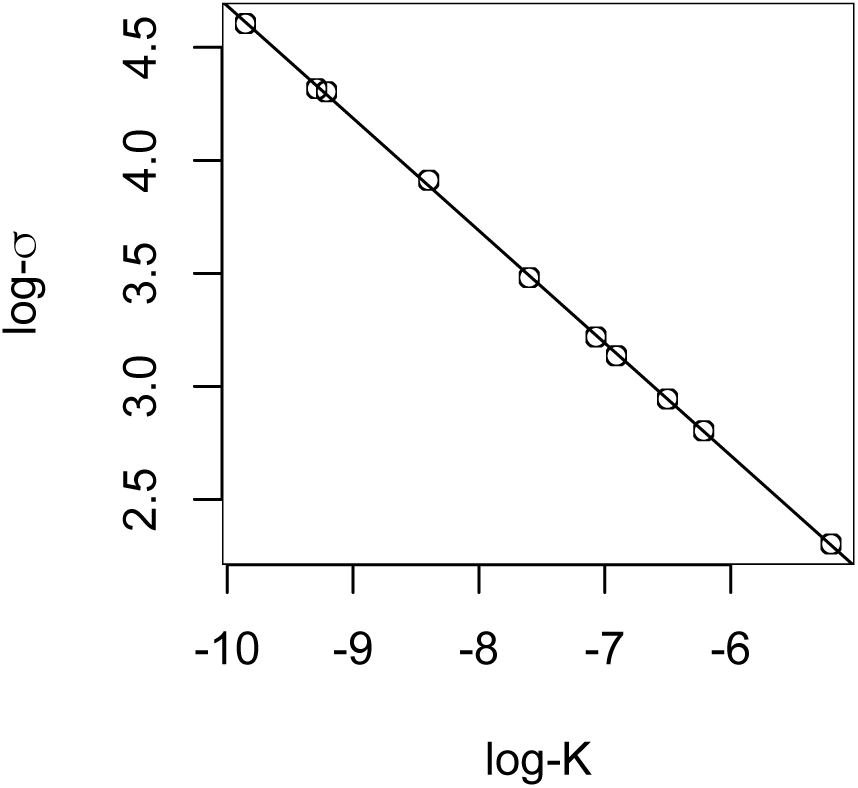
The minimum dispersal rate (*σ*) supporting a stable population for different carrying capacity (*K*) values.

**Figure S4:**
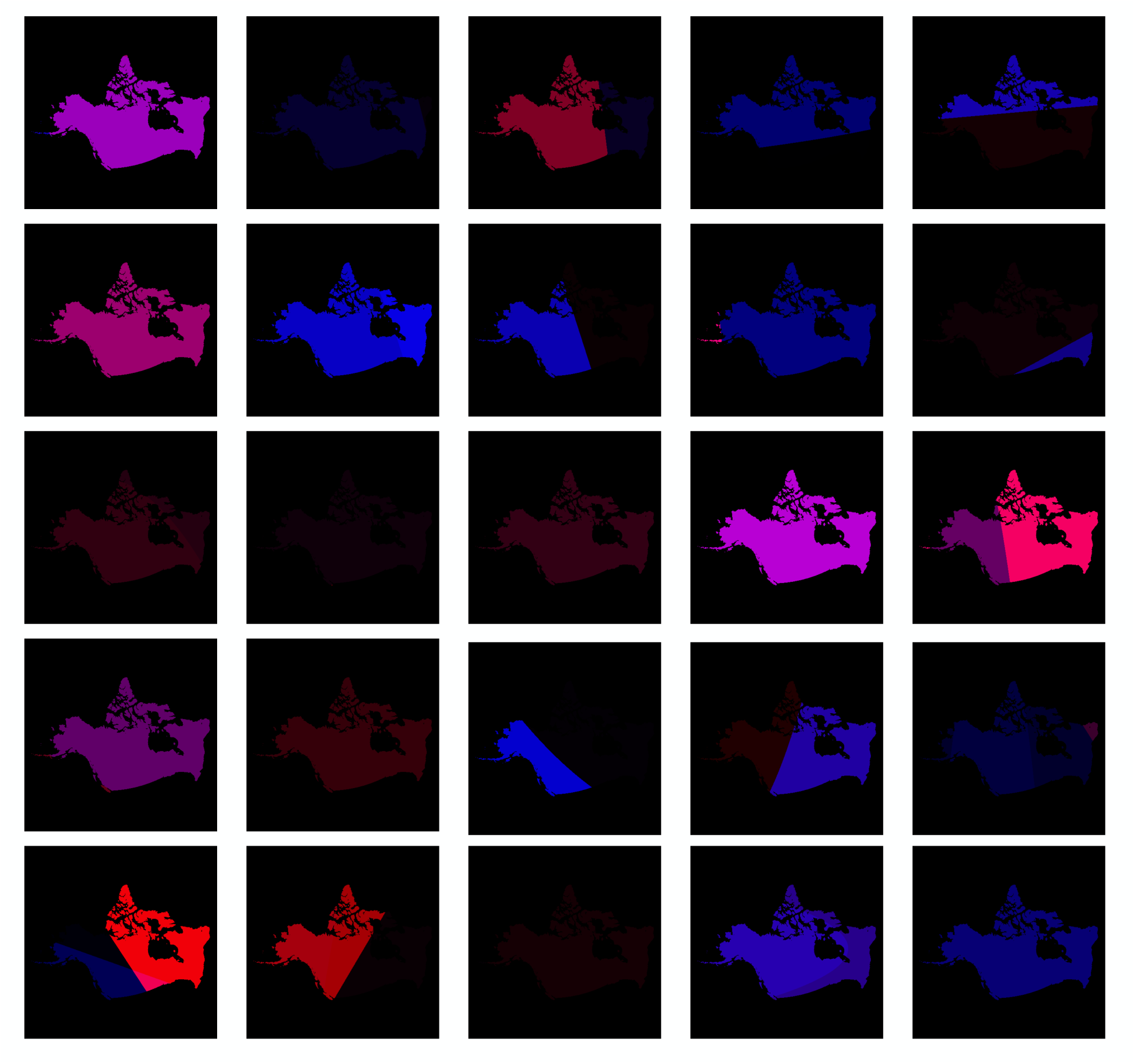
PNG renderings for a random selection of training maps for the North American grey wolf analysis. The blue channel conveys dispersal rate and the red channel conveys carrying capacity.

**Figure S5:**
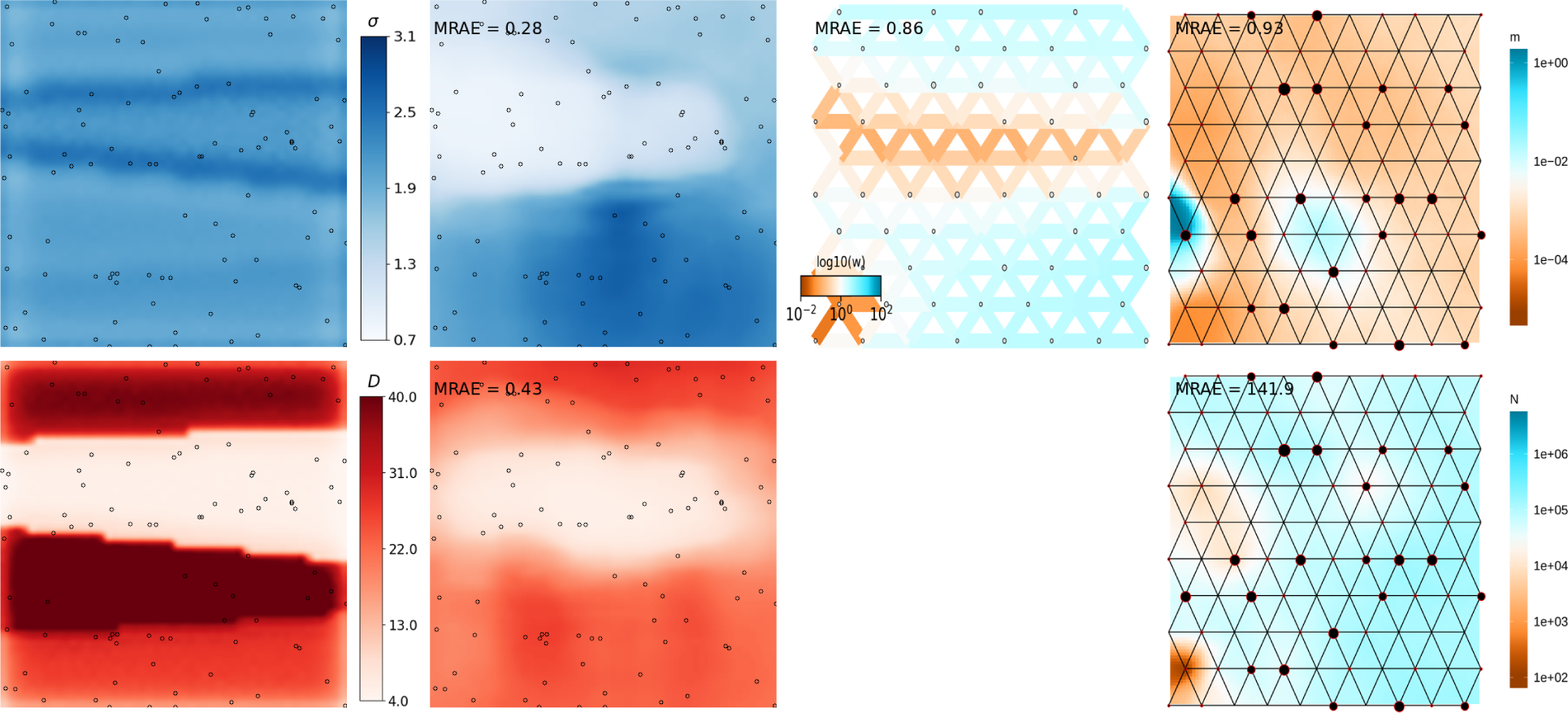
Predicted maps for a randomly selected, simulated test dataset. The leftmost column shows the ground truth maps for dispersal (top row) and density (bottom row). Columns 2-4 show estimated maps using three different methods: `mapNN`, `FEEMS`, and `MAPS` (respectively).

**Figure S6:**
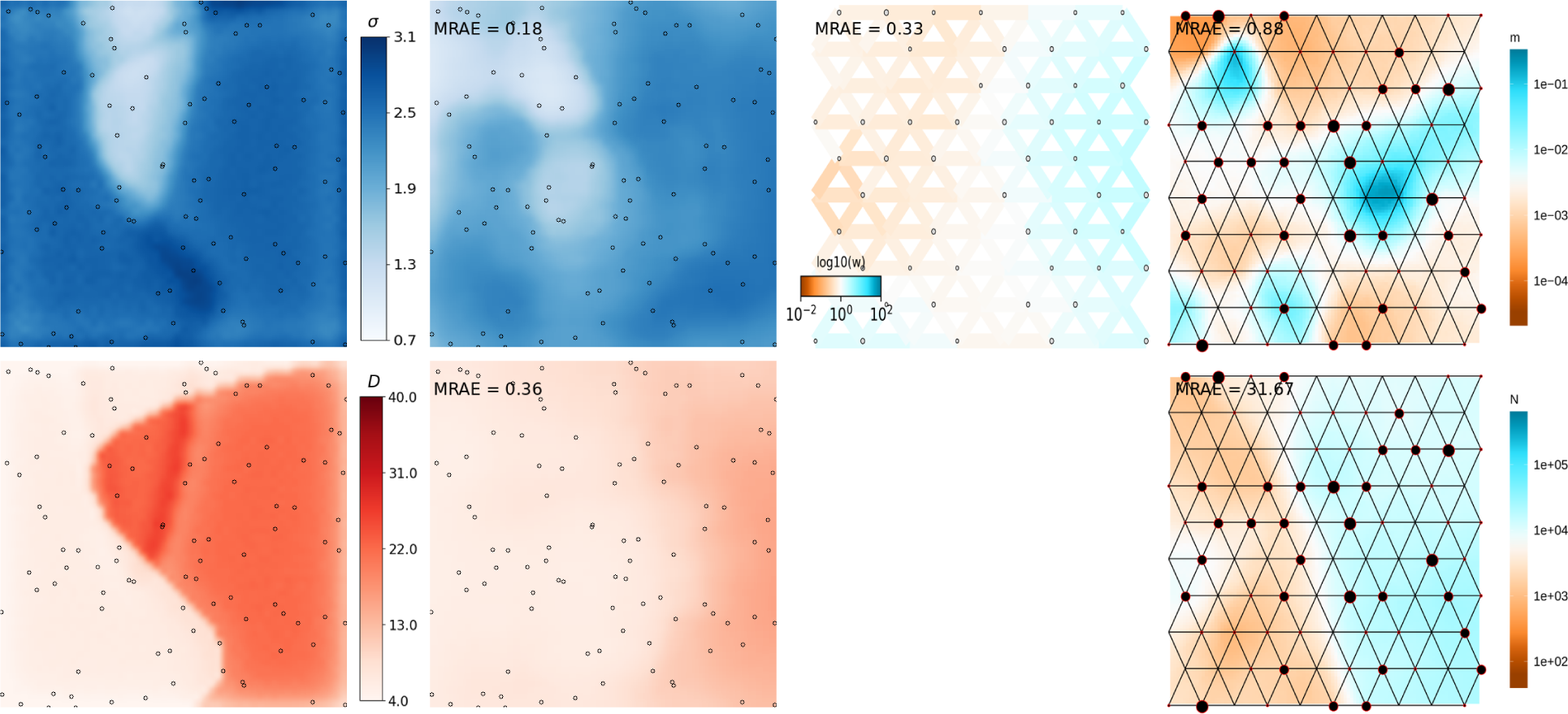
Predicted maps for a randomly selected, simulated test dataset. The leftmost column shows the ground truth maps for dispersal (top row) and density (bottom row). Columns 2-4 show estimated maps using three different methods: `mapNN`, `FEEMS`, and `MAPS` (respectively).

**Figure S7:**
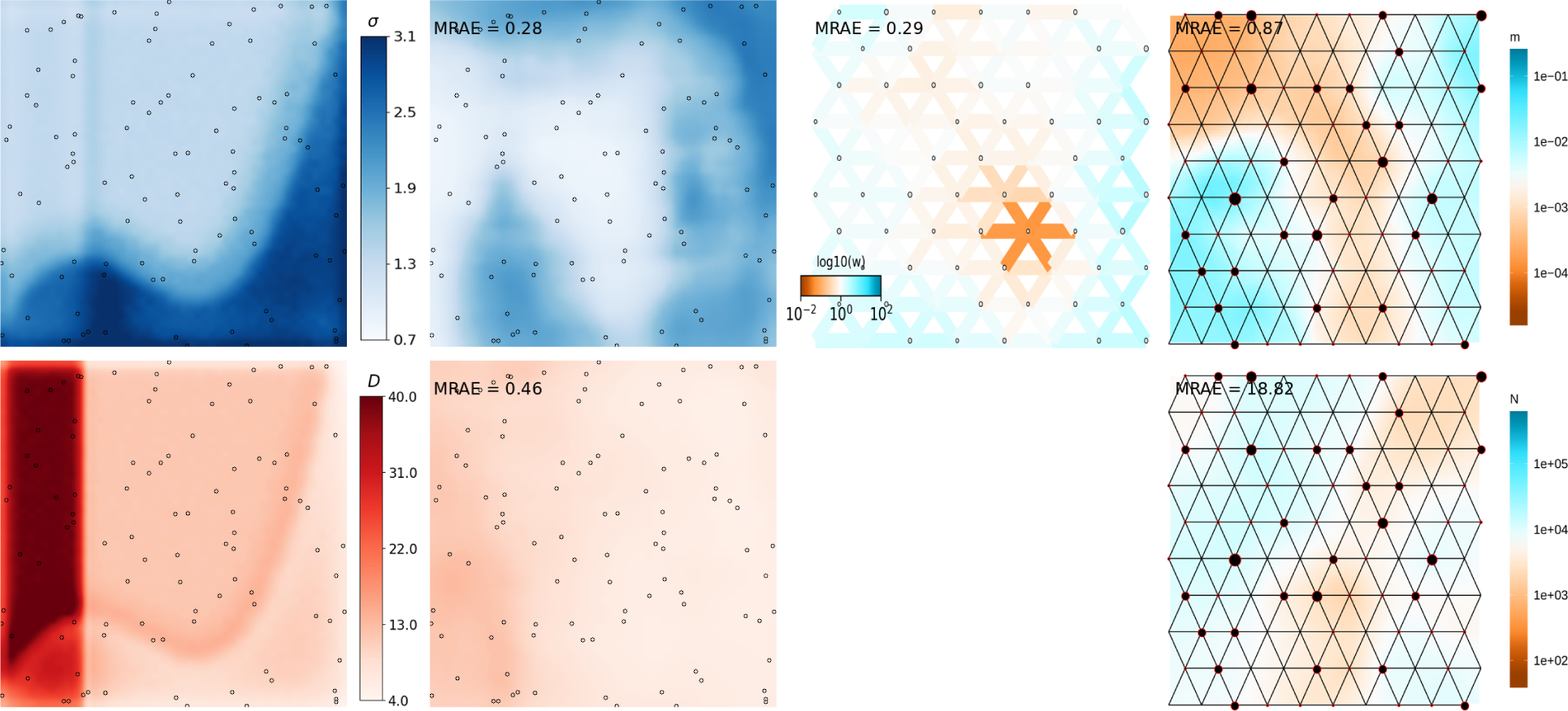
Predicted maps for a randomly selected, simulated test dataset. The leftmost column shows the ground truth maps for dispersal (top row) and density (bottom row). Columns 2-4 show estimated maps using three different methods: `mapNN`, `FEEMS`, and `MAPS` (respectively).

**Figure S8:**
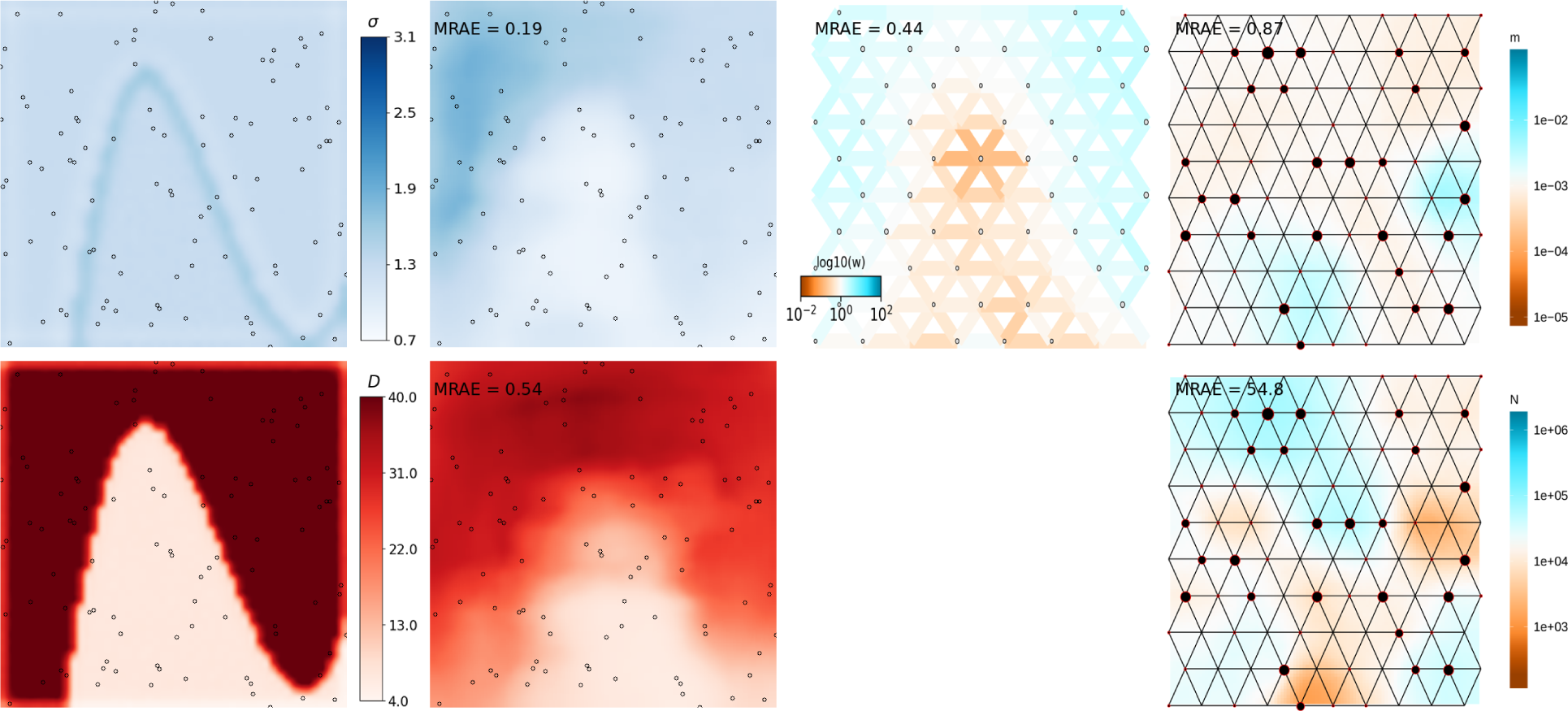
Predicted maps for a randomly selected, simulated test dataset. The leftmost column shows the ground truth maps for dispersal (top row) and density (bottom row). Columns 2-4 show estimated maps using three different methods: `mapNN`, `FEEMS`, and `MAPS` (respectively).

**Figure S9:**
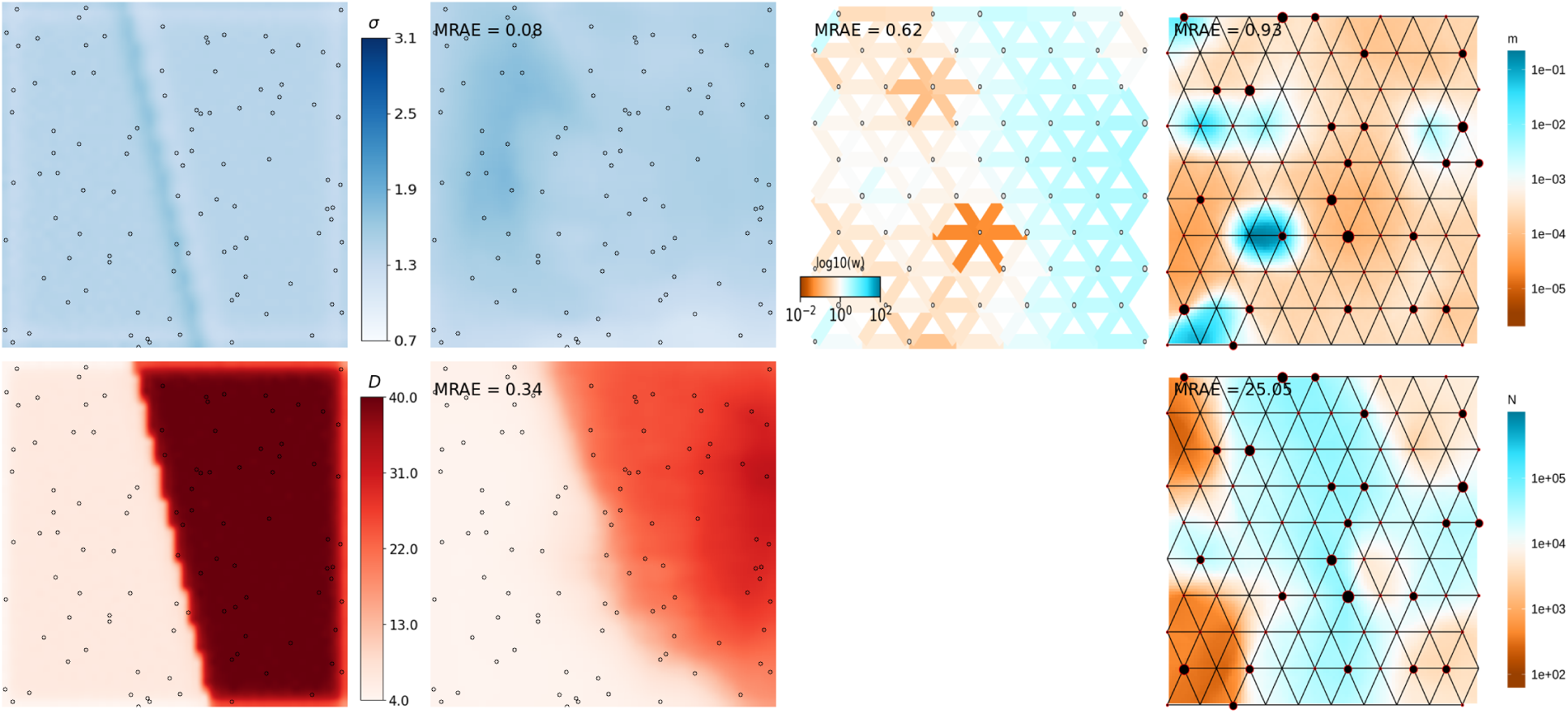
Predicted maps for a randomly selected, simulated test dataset. The leftmost column shows the ground truth maps for dispersal (top row) and density (bottom row). Columns 2-4 show estimated maps using three different methods: `mapNN`, `FEEMS`, and `MAPS` (respectively).

**Figure S10:**
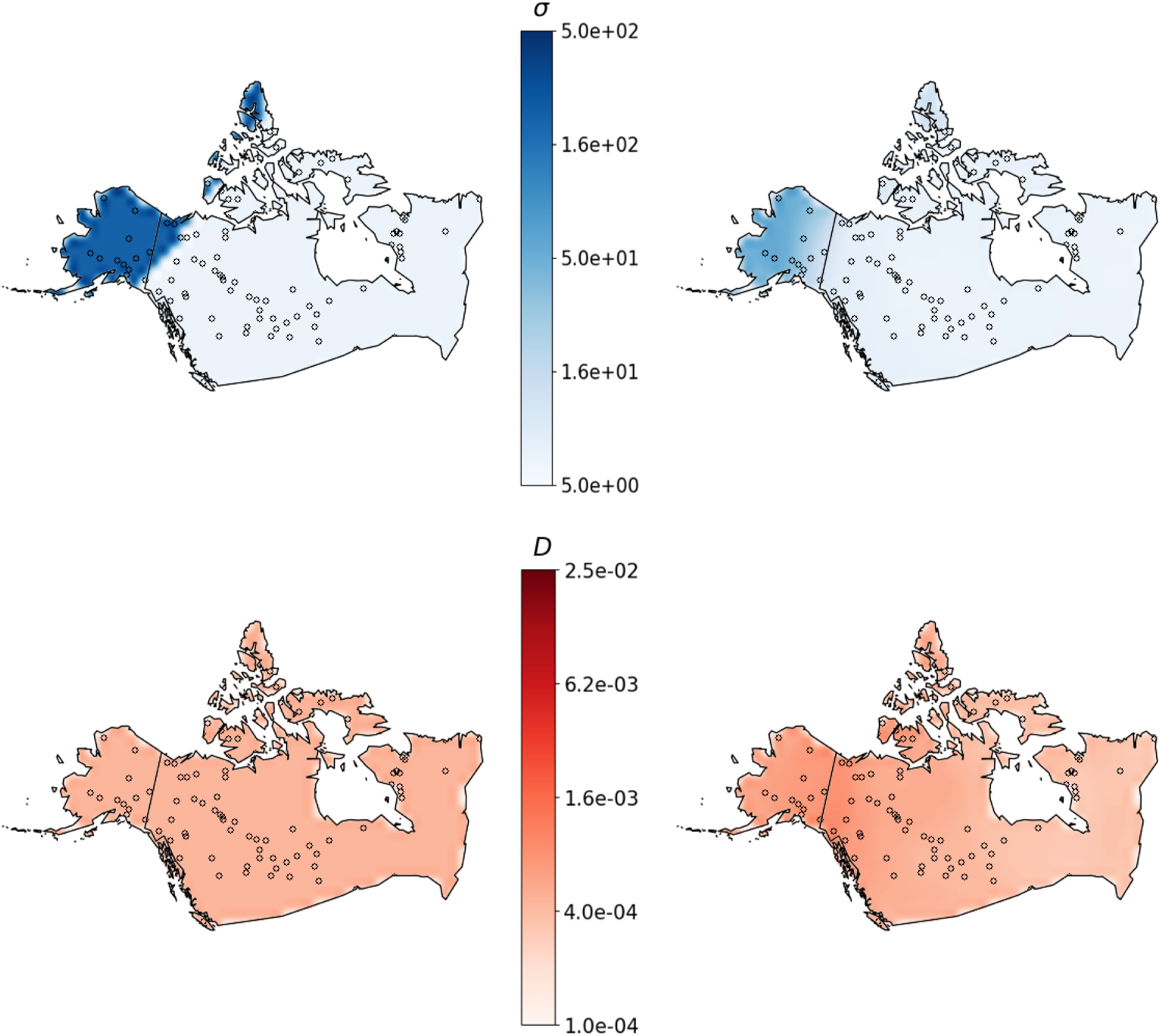
Predicted maps for a randomly selected, simulated test dataset for the North American grey wolf analysis. The left column shows the ground truth maps for dispersal (top row) and density (bottom row). The right-hand column shows estimated maps from `mapNN`.

**Figure S11:**
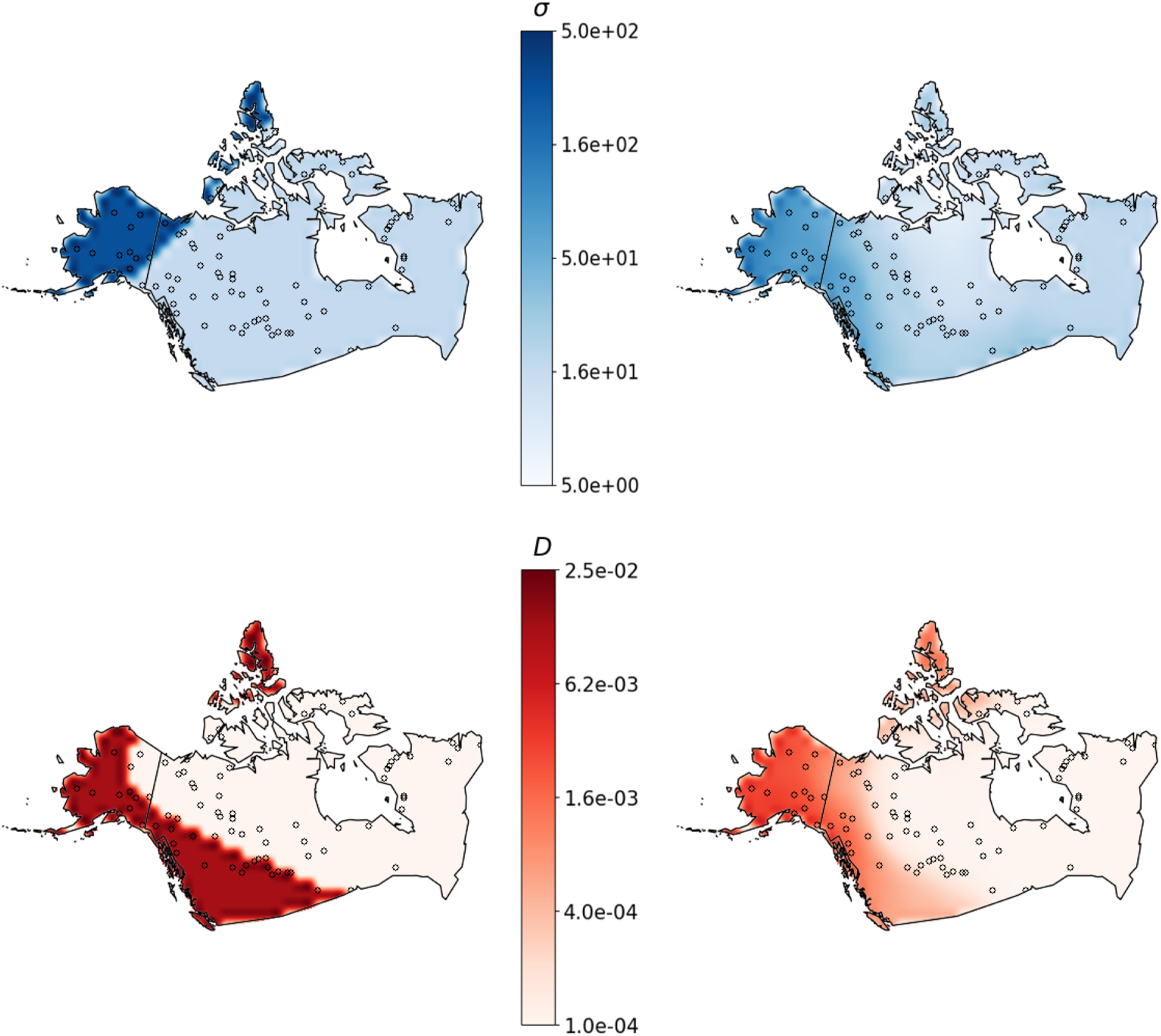
Predicted maps for a randomly selected, simulated test dataset for the North American grey wolf analysis. The left column shows the ground truth maps for dispersal (top row) and density (bottom row). The right-hand column shows estimated maps from `mapNN`.

**Figure S12:**
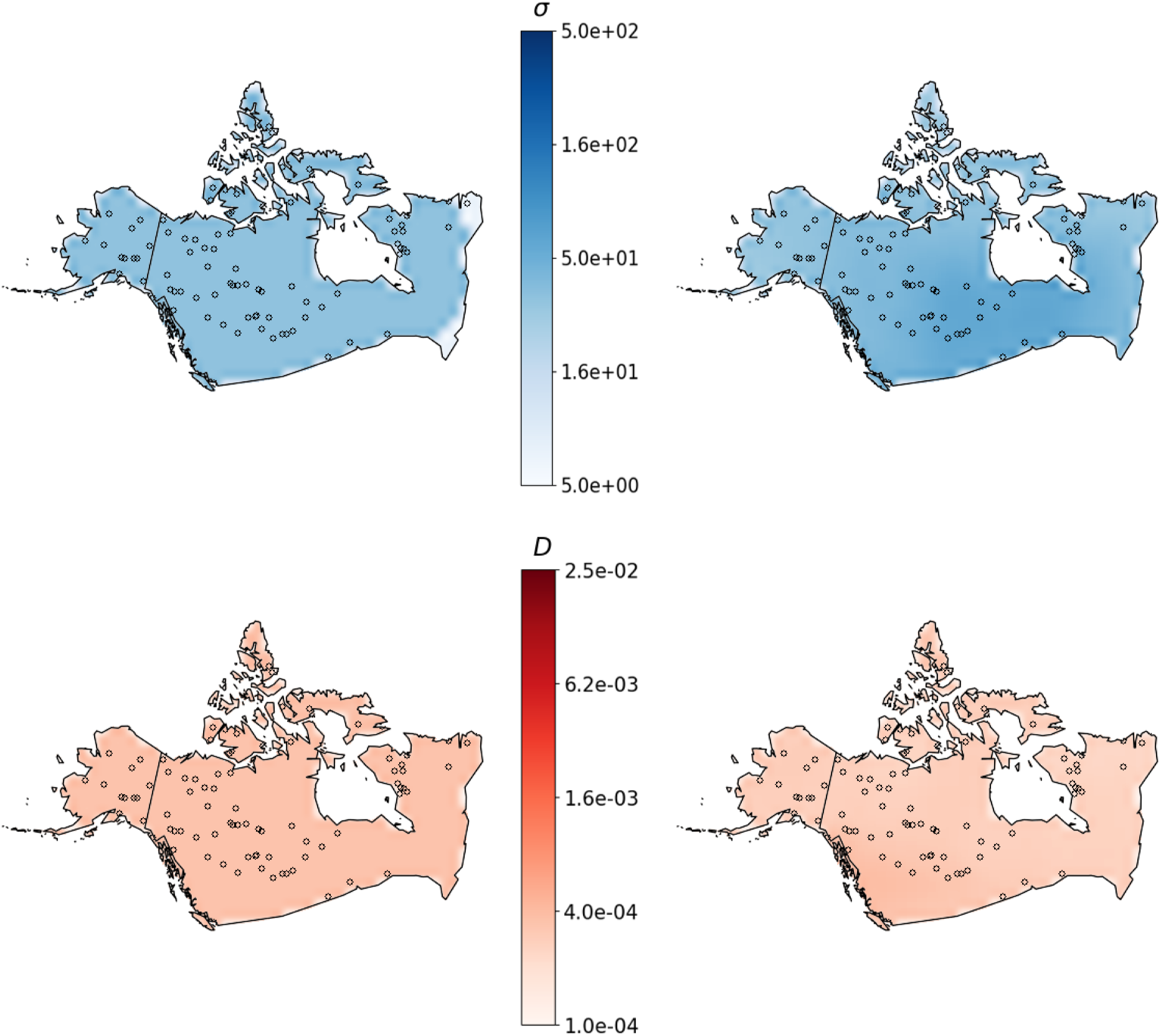
Predicted maps for a randomly selected, simulated test dataset for the North American grey wolf analysis. The left column shows the ground truth maps for dispersal (top row) and density (bottom row). The right-hand column shows estimated maps from `mapNN`.

**Figure S13:**
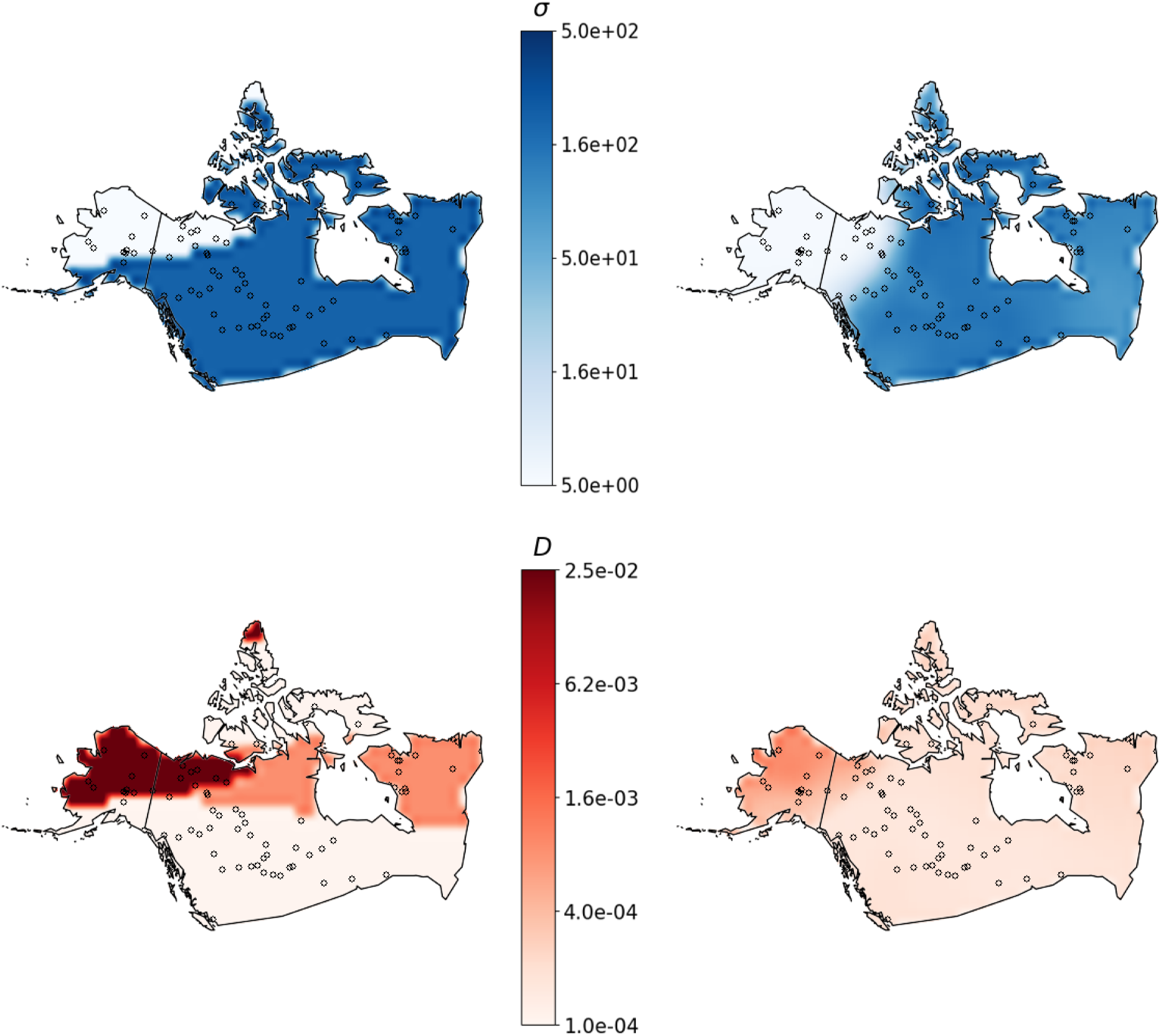
Predicted maps for a randomly selected, simulated test dataset for the North American grey wolf analysis. The left column shows the ground truth maps for dispersal (top row) and density (bottom row). The right-hand column shows estimated maps from `mapNN`.

**Figure S14:**
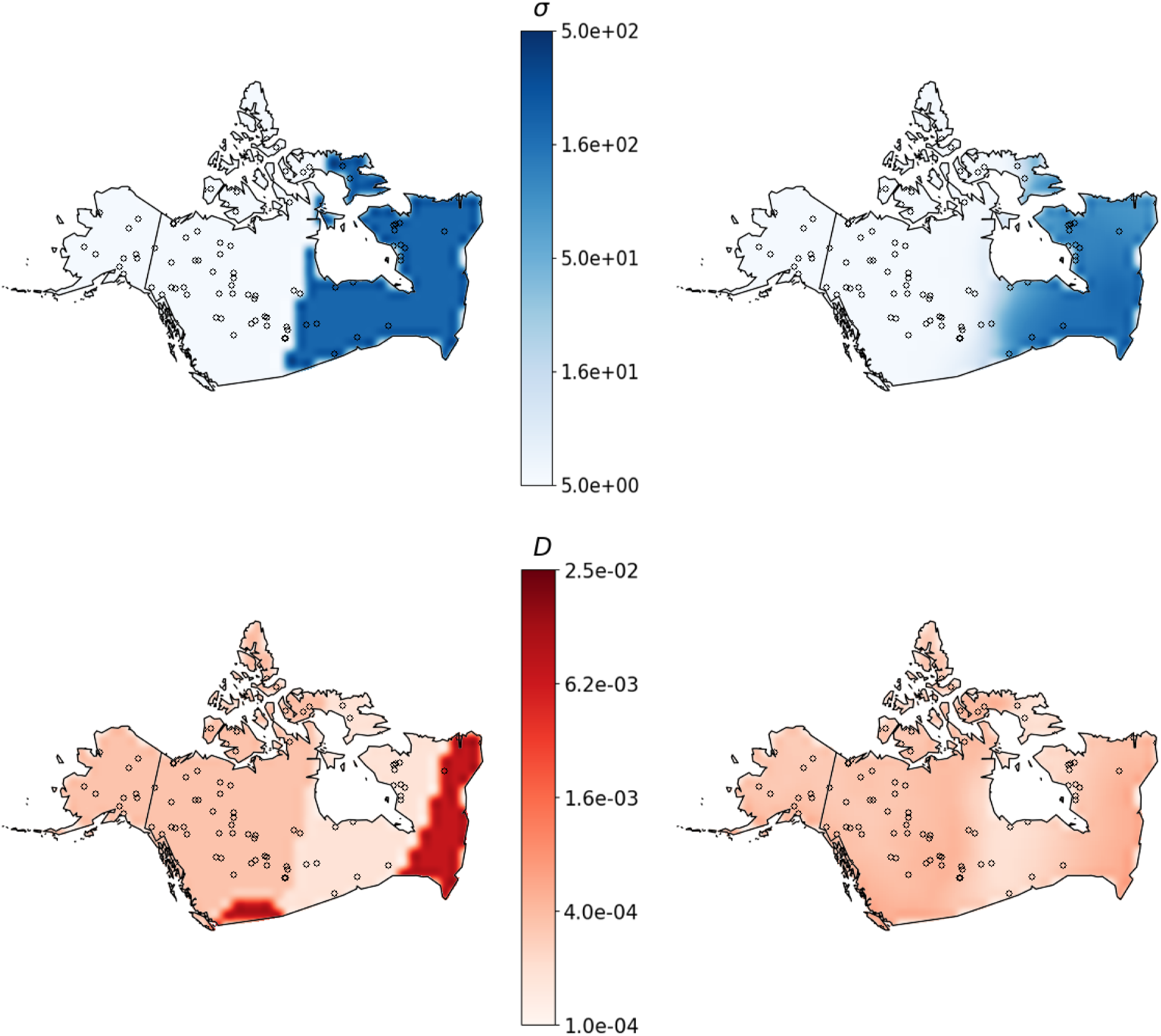
Predicted maps for a randomly selected, simulated test dataset for the North American grey wolf analysis. The left column shows the ground truth maps for dispersal (top row) and density (bottom row). The right-hand column shows estimated maps from `mapNN`.

**Figure S15:**
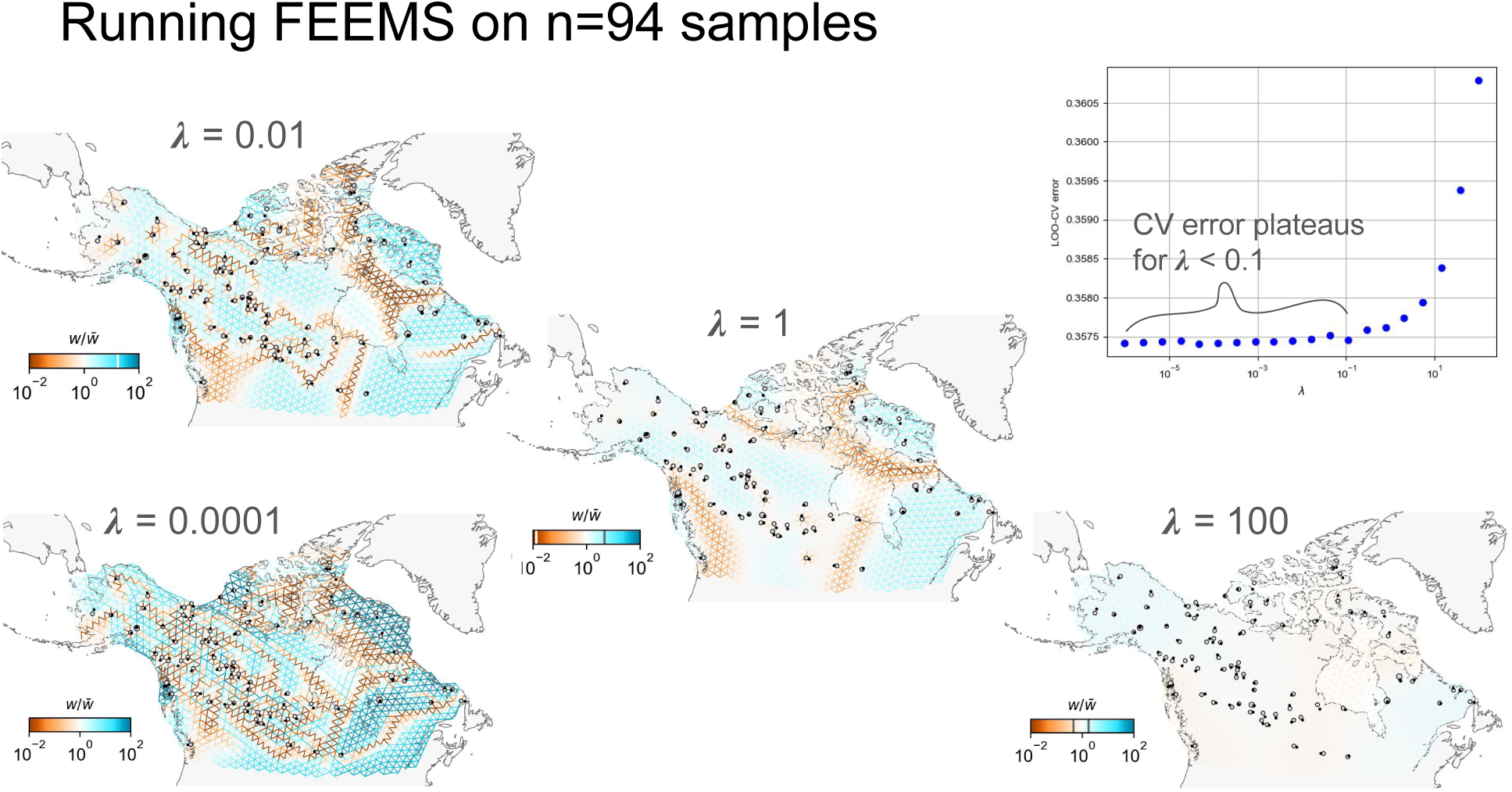
FEEMS run on the same *n* = 94 individuals analyzed in the current study.

